# EXPLANA: A user-friendly workflow for EXPLoratory ANAlysis and feature selection in cross-sectional and longitudinal microbiome studies

**DOI:** 10.1101/2024.03.20.585968

**Authors:** Jennifer Fouquier, Maggie Stanislawski, John O’Connor, Ashley Scadden, Catherine Lozupone

## Abstract

**Motivation:** Longitudinal microbiome studies (LMS) are increasingly common but have analytic challenges including non-independent data requiring mixed-effects models and large amounts of data that motivate exploratory analysis to identify factors related to outcome variables. Although change analysis (i.e. calculating deltas between values at different timepoints) can be powerful, how to best conduct these analyses is not always clear. For example, observational LMS measurements show natural fluctuations, so baseline might not be a reference of primary interest; whereas, for interventional LMS, baseline is a key reference point, often indicating the start of treatment.

**Results:** To address these challenges, we developed a feature selection workflow for cross-sectional and LMS that supports numerical and categorical data called EXPLANA (EXPLoratory ANAlysis). Machine-learning methods were combined with different types of change calculations and downstream interpretation methods to identify statistically meaningful variables and explain their relationship to outcomes. EXPLANA generates an interactive report that textually and graphically summarizes methods and results. EXPLANA had good performance on simulated data, with an average area under the curve (AUC) of 0.91 (range: 0.79-1.0, SD = 0.05), outperformed an existing tool (AUC: 0.95 vs. 0.56), and identified novel order-dependent categorical feature changes. EXPLANA is broadly applicable and simplifies analytics for identifying features related to outcomes of interest.

## Background/Introduction

Currently, scientific studies often include a collection of complex multiomic data,^1^ such as microbiome,^2^ transcriptome,^3^ and metabolome,^4^ and it is of interest to explore whether any novel features, or collections of features, may be related to an outcome variable. Adding to the complexity, researchers often collect other data from individuals that may impact an outcome, such as demographic and health data, or surveys on diet or medications. The growing quantity of available data complicates statistical decisions regarding variable inclusion, which is often based on hypotheses that motivated initial study design. Additionally, studies can include both categorical and numerical variables and can often contain non-independent longitudinal data, posing greater statistical challenges. As research advancements are made, collaborative efforts with different research laboratories produce more data per study, and human biases are often introduced during study design and analytics. These challenges have ultimately stimulated a growing interest in data-driven methods.

One field particularly impacted by an abundance of data is microbiome research, which focuses on characterizing the community of viruses, fungi and bacteria and their genes. Characterization of the microbiome is often performed by 16S ribosomal RNA (rRNA) gene sequencing, which identifies the bacteria and archaea in an environment. One well-studied microbial environment is the gut microbiome because of the metabolic potential of the bacterial community and its association with numerous human diseases, including obesity,^5^ depression,^6^ autism,^7^ cancer,^8,9^ HIV^10^ and cardiovascular disease.^11^ The gut microbiome relationship to human disease suggests that gut microbiome modification through interventions like dietary changes, probiotics, or fecal microbial transplants may provide disease prevention or treatment options.

To understand changes in health outcomes and to address the impact of individual variation, longitudinal studies that collect data from multiple individuals, at different timepoints, are essential. In addition to these studies often containing diverse subject data (with numerical and categorical variables), they include repeated measurements on individuals which requires special statistical consideration to identify relationships between features within non-independent data.^12^ Random Forest (RF)^13^ based machine learning (ML) approaches are powerful for combining different data types to predict outcomes and identify important features. RFs work well with high-dimensional data (more features than samples/instances),^14^ find non-linear relationships, work with non-normal data distributions, and are more interpretable than many other ML models because they are based on simple decision trees. Implementation of interpretable methods can improve accessibility of complex tools. Additionally, mixed-effects RF (MERF)^15^ models can be used for longitudinal study designs. However, numerous challenges can hinder effective application of these methods.

MERFs can be run on original (raw) data from longitudinal studies or by using changes (Δs) between different reference timepoints, which can reveal unique insights in some studies.^16–20^ However, the research question of interest can affect decisions regarding optimal calculation of Δs. In some designs, such as interventions, or some observational studies with an expected trend over time (e.g., gut microbiome changes over the first years of a baby’s life^16^), changes are expected to be compared to a baseline value, so Δs can be calculated using baseline as a reference.^17,18^ However, some observational studies have no meaningful baseline, and it might be of interest to relate an outcome variable to changes in predictors between adjacent timepoints or all pairs of timepoints.^21,22^ For instance, in an observational longitudinal study of children with autism spectrum disorder (ASD) that we conducted,^22^ children with ASD were evaluated over time to identify relationships between diet, gastrointestinal distress, or the microbiome and ASD-associated behaviors. Because of high interpersonal gut microbiome variation, this LMS revealed relationships between the gut microbiome and ASD behaviors as a correlation between the degree of microbiome change and ASD behavior change between timepoints. However, because more than two timepoints were studied, and because the first timepoint was not a meaningful baseline, pairwise analyses were performed. Pairwise analysis is useful for identification of effects that are time-delayed (i.e., a change from time 2 to time 4), order-dependent, or reference dependent. Different longitudinal study designs highlight the importance of understanding differences/changes (Δs) in features, for each subject over time, and that feature changes differ depending on their reference values. Statistical methods differ regarding how and when to apply change analysis and can even lead to different conclusions.^23^ Thus, our method compares results from original and Δ datasets for a more complete picture of a longitudinal study.

Another analytic challenge encountered in the application of RFs to complex microbiome studies is the integration of microbiome data as predictor variables (i.e. features) with other data types (e.g., surveys or clinical reports with numerical and categorical data). To the best of our knowledge, there are no software tools that create and select order-dependent categorical feature changes that impact an outcome variable. For example, the drugs amiodarone and quinidine for heart arrhythmia treatment have an interaction that could lead to a dangerously rapid heartbeat,^24^ but an interaction risk is higher if amiodarone precedes quinidine since amiodarone has a much longer elimination half-life (days^25^ vs hours^26^). This example highlights how calculating order-dependent categorical Δs might uncover relationships that have differential impact if introduced in opposite order, such as in crossover study designs (AB/BA designs). This led us to the hypothesis that unique features dependent on different contexts of change could be identified, including novel order-dependent categorical features by tracking text changes as an engineered feature value (e.g., “amiodarone quinidine”).

Finally, another key challenge is performing complex LMS analytics in a reproducible way that facilitates communication about results. These workflows can involve inputs of diverse data types, calculation of Δs with different reference points, feature selection using mixed-effects ML methods, and methods for explaining why features were selected, in addition to their importance ranks. Although there are tools for feature selection in microbiome data,^16,27–31^ none provide the combination of methods described here. For example, timeOmics^31^ is useful for multi-omic integration with an emphasis on time as the outcome, while other goals are to identify features related to different outcome variables over time. QIIME 2 longitudinal feature-volatility,^16^ a feature selection tool provided as part of a very popular microbiome analytics platform, allows for looking at different outcomes, but does not incorporate metrics that explain the selected feature’s impact on an outcome. Both tools, although useful for longitudinal analytics, do not incorporate categorical Δs. Although individual tools for data-pre-processing, application of RFs or MERFs, and downstream interpretation of results also exist, it is cumbersome for scientists to research and implement this complex array of tools.

For these reasons, we developed EXPLANA (EXPLoratory ANAlysis), a data-driven feature selection workflow that streamlines hypothesis generation, accommodating longitudinal and cross-sectional data, as well as both numerical and categorical variables. EXPLANA can identify unique features important in different contexts of change, including order-dependent categorical features related to changes in outcomes. The combination of novel and existing methods to address analytic challenges provides broad applicability for scientific discovery.

## Results

### Software Workflow Summary

EXPLANA was developed to create a comprehensive feature selection report. The workflow is guided by directions from a configuration file where the user provides dataset paths, selects optional preprocessing steps per dataset, and defines inputs and the outcome of interest (Figure 1). The input datasets are merged to form the comprehensive *Original* dataset. If more than one timepoint is sampled per subject, feature changes are calculated using different reference points to uncover important features in different contexts of change. Thus, the *Original* dataset is used to compute three Δ datasets, *First*, *Previous*, and *Pairwise* (Figure 2). Differences are calculated as follows: For *First*, compared to baseline/first measures. For *Previous*, compared to the immediately previous timepoint. For *Pairwise*, all pairwise comparisons between timepoints. Notation throughout is as follows: Timepoints 1, 2, 3, etc. are referred to as T1, T2, T3, etc., respectively. Accordingly, the difference between T1 and T2 is T1_T2, and labeled in chronological order. For T1_T2, T1 is the reference and is subtracted from T2 (Figure 2).

**Figure 1.**
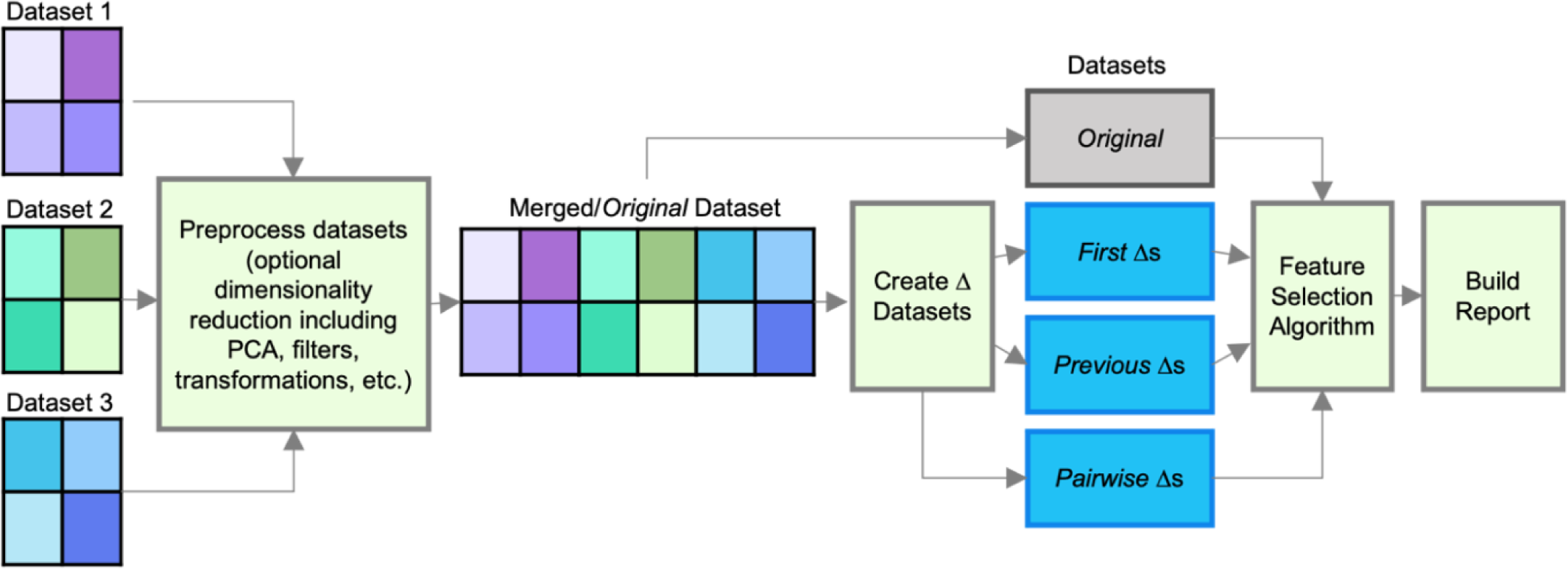
Feature selection workflow diagram. Individual datasets can be preprocessed separately to reduce dimensionality using a variety of methods, including principal components analysis (PCA), center-log-ratio (CLR) transformation or filters. Datasets are merged to form the *Original* dataset prior to creation of Δ datasets for longitudinal studies. *First, Previous,* and *Pairwise* Δ datasets, as shown in blue, are created as explained in Figure 2. During Δ dataset creation, distance matrices can be incorporated. Feature selection is performed for up to four models built from each dataset (*Original, First, Previous,* and *Pairwise*). An .html report is created which summarizes features selected per model.

**Figure 2.**
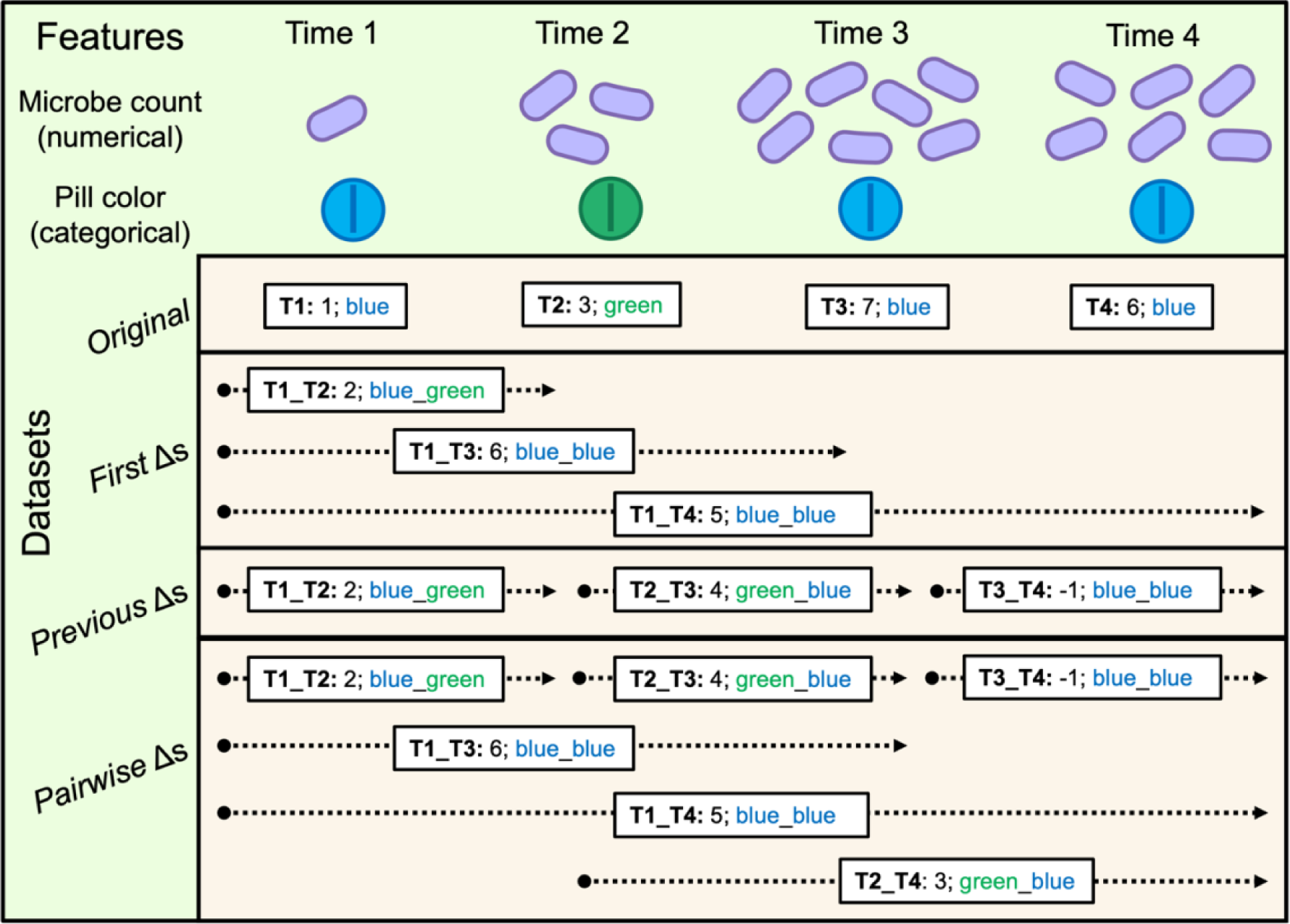
Example calculations of *First, Previous* and *Pairwise* Δ datasets for numerical and categorical features in a four timepoint study. The *Original* dataset includes original feature values (without change analysis) and Δ datasets contain differences/changes in features, per subject, between timepoints. Different reference points are used for each Δ dataset. For categorical features (e.g. pill color), text is used to track the order of categorical changes and for numerical features (e.g. microbe count), the reference is subtracted from the later timepoint. The comparison between two timepoints is indicated before each colon (e.g., a Δ between timepoint 1 and 2 is indicated as T1_T2). For a four timepoint study: *First* Δs: T1_T2, T1_T3, T1_T4; *Previous* Δs: T1_T2, T2_T3, and T3_T4; and *Pairwise* Δs: T1_T2, T2_T3, T3_T4, T1_T3, T1_T4, and T2_T4.

EXPLANA accommodates numerical and categorical predictor and outcome variables, including novel functionality to track categorical feature changes over time. Microbiome-specific challenges addressed include the option to use a center-log-ratio (CLR) transformation for compositional data,^32^ as well as incorporating distance matrices during Δ calculations, allowing users to evaluate differences between microbiome samples, such as those calculated with UniFrac or other beta diversity measures.^33^ For feature selection, MERFs are used as the ML method for independent, repeated measures data, otherwise RFs are used. The Boruta^34^ method combined with SHAP^35^ (BorutaSHAP^36^) is used to rank features by their importance for model performance, identify which contribute to a more accurate prediction of the outcome than expected by random chance, and produce plots to assess whether features have a positive or negative impact on an outcome, thereby improving results interpretation.

Upon workflow completion, a report is generated that contains a description of the analysis, as well as tables and figures that explain why features were selected (Figure 3). The ease of running different datasets through the EXPLANA workflow enables easier exploratory analyses of datasets, such as performing stratified analyses. For example, to do stratified analyses, the user simply needs to use small R scripts inside the configuration file which will be documented in the report.

**Figure 3.**
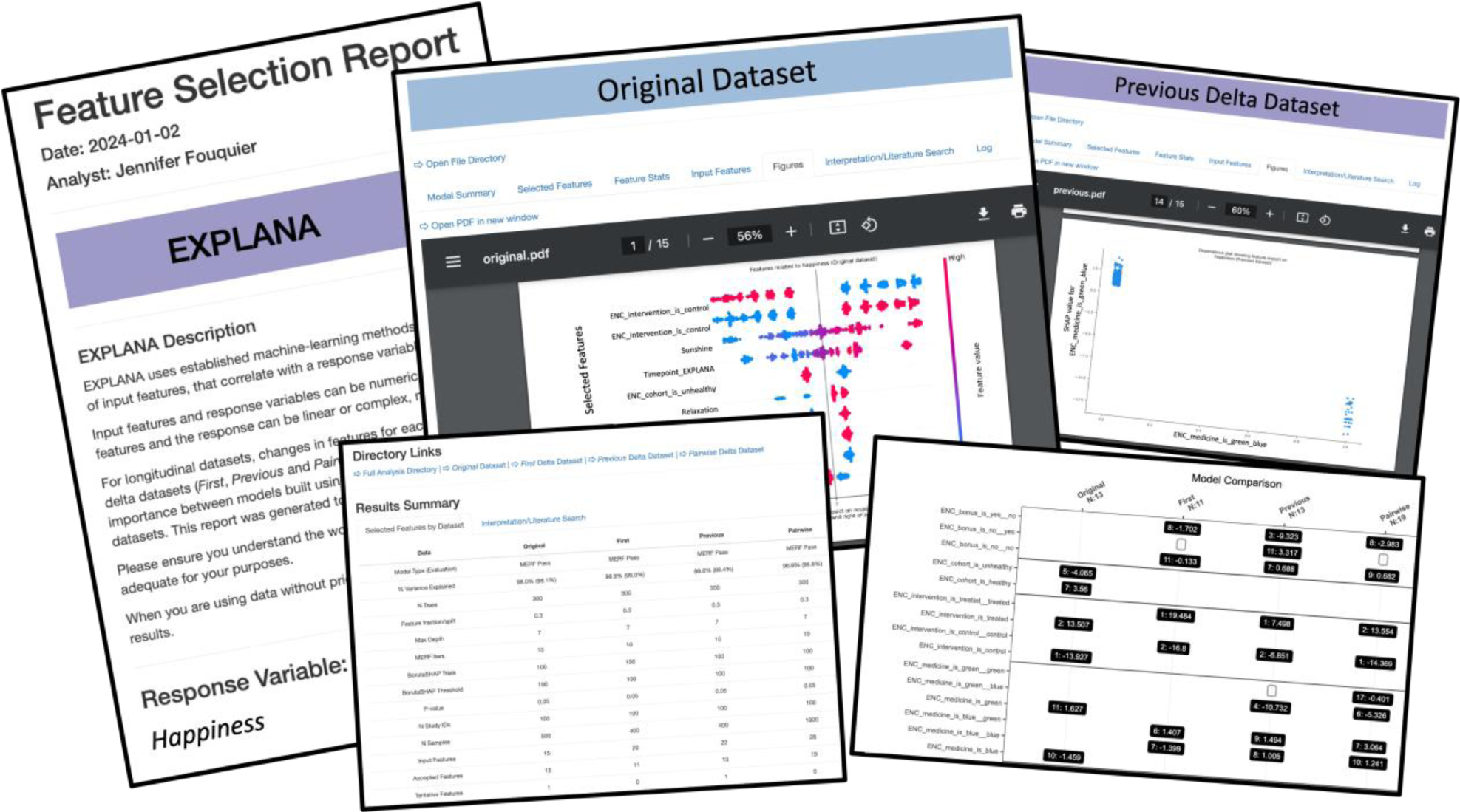
Screenshots from a feature selection report. The report is in an interactive .html format to facilitate interpretation of complex machine-learning method results through figures and written explanations. The report includes a written summary of methods containing arguments and information from the configuration file about the analytic decisions. The dynamic methods can be copy/pasted into manuscripts for efficiency; tables and figures summarize feature selection results for models built from *Original* and, if longitudinal, *First*, *Previous* and *Pairwise* Δ datasets. Figures include SHAP summary plots and SHAP dependence plots. Links are provided to directories containing files used for report generation to facilitate exploration of data and results.

### Workflow evaluation and feature selection using a simulation study

The workflow was evaluated using simulation studies and published datasets (both detailed in *Methods*). A simulation study on longitudinal happiness, called *SimFeatures*, was created for performance evaluation and modeled as an intervention including 100 individuals treated with one of two therapies to improve happiness over five timepoints. Happiness is based on a numerical score where higher values indicate better mental health. For interpretability, features are recognizable as factors that could affect real-life happiness such as relaxation, sunshine, salary, medication, etc. Categorical and numerical features were included with and without relationships to happiness, thus representing predictive and not predictive engineered features (Supplemental Table 1). Some features were designed to be important only in some of the four models (Figure 2; *Original, First, Previous* or *Pairwise*) to validate whether the tool could select unique features dependent on different contexts of change.

A simulated microbiome feature table (compositional data) was also created using MicrobiomeDASim^37^ with 25 differentially abundant microbes linearly correlated to happiness changes over time and 175 that are not related. The dataset with the outcome variable and simulated microbes was called *SimMicrobiome*. The dataset with *SimFeatures* and *SimMicrobiome* combined was called *SimFeaturesMicrobiome0.* To evaluate the effects of including many features without a relationship to the outcome, an increasing number of random variables from a variety of data distributions were added to the *SimFeaturesMicrobiome* dataset. The number of random variables is indicated in the dataset names. Thus, the five simulation studies used for workflow evaluation are *SimFeatures*, *SimMicrobiome*, *SimFeaturesMicrobiome0*, *SimFeaturesMicrobiome500*, and *SimFeaturesMicrobiome1000*. (Supplemental Table 1).

Workflow evaluation was performed by appropriate selection or rejection of engineered features. True positives (*selected* predictive features; TPs) and true negatives (*rejected* not-predictive features; TNs) were considered correctly classified, while false positives (*selected* not-predictive features; FPs) and false negatives (*rejected* predictive features; FNs) were considered incorrectly classified (Supplemental Figure 1). These datasets allowed us to: 1) evaluate workflow performance from classification accuracy of engineered features and 2) to test the hypothesis that unique features dependent on different contexts of change could be identified, including novel order-dependent categorical changes related to an outcome.

EXPLANA was used to select and rank features related to happiness for the five simulation studies for all models (*Original, First, Previous* or *Pairwise*) to evaluate performance (Figure 4, Table 1, Supplemental Figure 2). Area under the curve (AUC) and F1-score (a metric that accounts for both precision and recall; (Supplemental Figure 1) respective ranges for *Original, First, Previous* and *Pairwise* were: 0.79-1.00 and 0.83-1.00 *SimFeatures*; 0.80-0.96, and 0.73-0.87 for *SimMicrobiome*; 0.88-0.94 and 0.78-0.91 for *SimFeaturesMicrobiome0*; 0.87-0.95 and 0.69-0.91 for *SimFeaturesMicrobiome500*; and 0.90-0.95 and 0.66-0.92 for *SimFeaturesMicrobiome1000. Original* yielded the highest F1-scores and AUCs. The average workflow ability to recall predictive features was good/excellent (average 0.87, SD = 0.09) and good for precision (avg 0.82, SD = 0.14), with some Δ datasets having lower precision or recall. Of the four models analyzed with EXPLANA for *SimMicrobiome*, *Previous* had the lowest percent variation explained (65.4%), a low recall (0.60), and failed to correctly classify some predictive features (Table 1). For simulated datasets with not-predictive microbes or random variables, *Pairwise* had the poorest precision. The lowest F1-score was for *SimFeaturesMicrobiome1000* using *Pairwise* (0.66), which had 30 FPs out of 1000 random variable that affected precision (0.55). However, the proportion of selected predictive features (recall) was good (0.81), and the AUC was 0.91 (see confusion matrix in Table 1).

**Figure 4.**
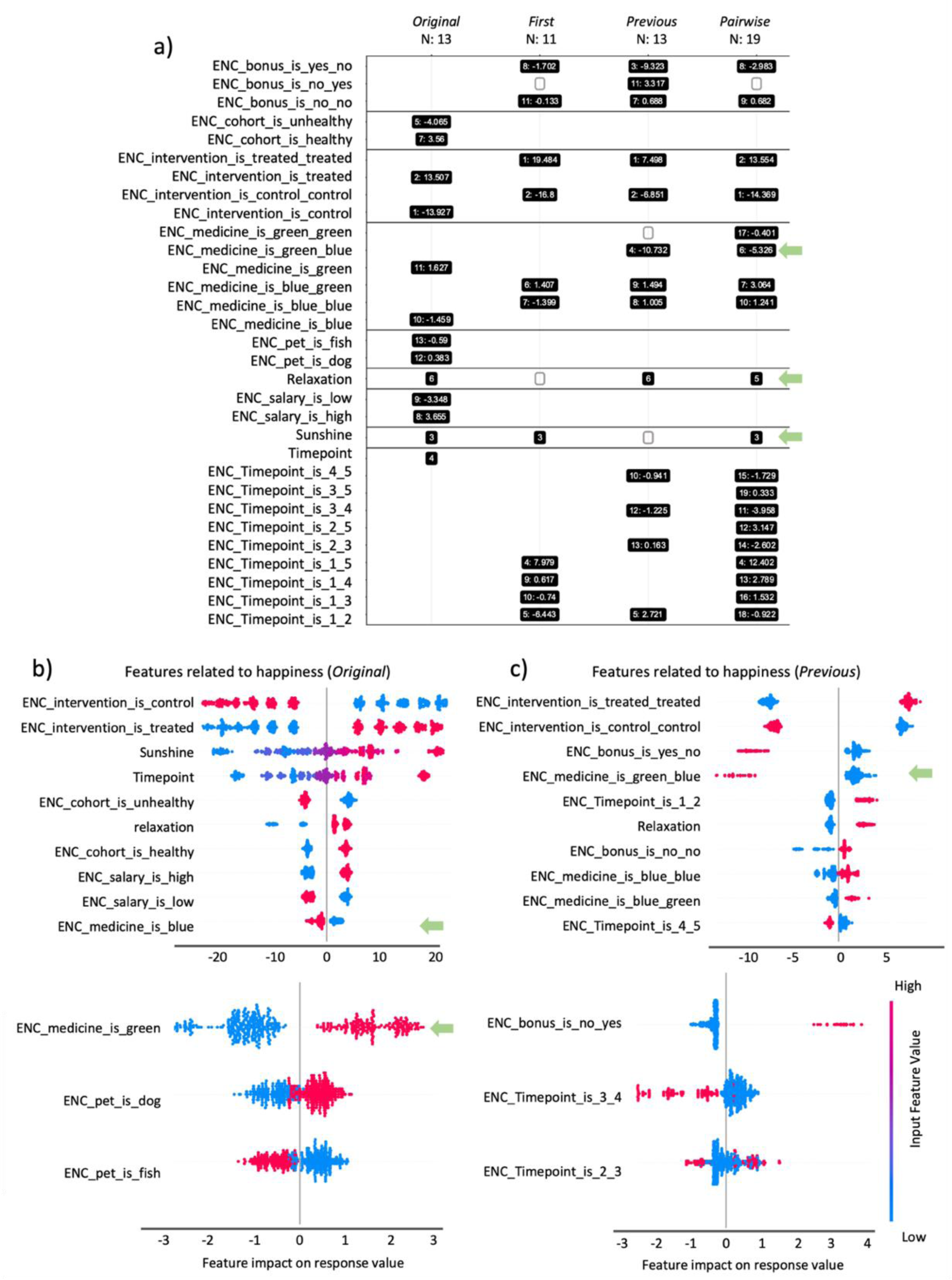
Feature selection report excerpts for one analysis using *SimFeatures* dataset. The *SimFeatures* dataset is a simulated longitudinal intervention with 100 individuals sampled over five timepoints (see Methods for detailed description). (a) Feature occurrence figure with selected features organized by model (*Original*, *First*, *Previous* and *Pairwise*) and ranked with one being the highest importance. For presence (yes/1) of a categorical variable, SHAP values are indicated following the colon. SHAP summary beeswarm plots for ranked selected features are shown for (b) *Original* and (c) *Previous* models. Each point represents one sample, and the horizontal position indicates impact on the outcome as indicated on the x-axis. Points to the left indicate a negative impact, and points to the right indicate a positive impact. The colors represent the selected feature values, where red is larger, and blue is smaller. For binary encoded features (‘ENC’) red is yes/1 and blue is no/0. Note that scales differ between the top and bottom SHAP plots, as they are grouped by a maximum of ten features per SHAP plot. Some features were designed to be identified in only certain models. Green arrows draw attention to key findings explained in *Results*. Interesting results include: ENC_medicine_is_green_blue (pill color), a categorical feature important in *Previous* and *Pairwise* models; “relaxation”, a numerical feature important in *Previous* and *Pairwise* and undetectable in *First* as described in the Discussion; and “sunshine” which was unable to be detected in *Previous* but a high rank in other models. Multiple analyses create a more comprehensive picture for longitudinal studies. 300 trees were used, with a feature fraction of 0.3, max depth of 7, with 10 iterations of mixed-effects Random Forests (MERFs), and 100 BorutaSHAP trials (100% importance threshold; p=0.05).

**Table 1.**
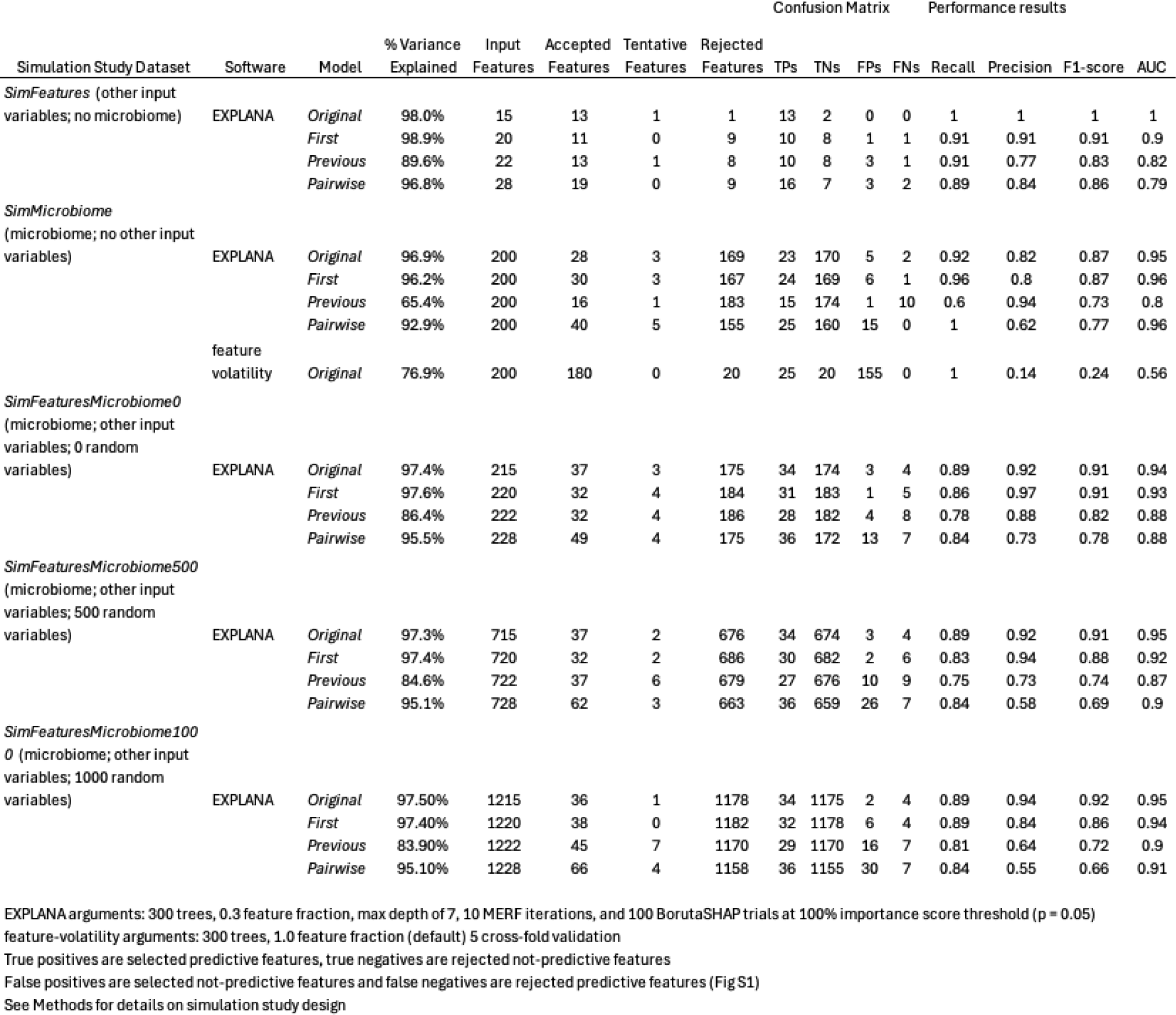
Performance Results from Classification Accuracy of Engineered Predictive and Not Predictive Features in Five Simulation Studies.

The features selected from the smallest dataset containing a variety of input feature types, *SimFeatures*, are shown in Figure 4a. Details about features and their motivation for workflow demonstration are explained in Supplemental Table 1. One feature that emphasizes the advantage of calculating Δs using the previous values or pairwise values rather than only first/baseline values is pill color, a categorical feature engineered to have a negative impact on happiness when blue pills were taken *after* green (“green_blue”), and not conversely. The feature green_blue had relatively high ranks and a negative impact on the outcome using *Pairwise* and *Previous* (respective rank:impact: *Pairwise* = 6/19:-5.3, and *Previous* 4/13: −10.7; Figure 4a and Figure 4c). Green was not a possible value at baseline/T1, therefore green_blue could not be identified with *First*. Green alone was selected with *Original* at a lower importance rank and a small positive impact (rank and impact: 11/13; +1.6) and blue was selected with a small negative impact (rank and impact: 10/13; −1.5). Green occurred at later timepoints, while happiness was also increasing, so selection of green with a small positive impact on happiness was appropriate in the *Original* despite being designed without independent effects. However, without the additional information provided by Δs, an assumption about positive impact on happiness when consuming green pills could have been made.

A numerical example that emphasizes the benefit of using different methods for calculating Δs in longitudinal analysis is “relaxation,” which was selected in *Original*, *Previous* and *Pairwise* models but not *First*. Relaxation had one value for baseline/T1 and a different equivalent value at all later timepoints, resulting in T1 comparisons to T2, T3, T4 and T5 having identical values (e.g., T1=1, T2-T5 = 5; differences compared to baseline would be 4 in *First*). The lack of change in *First* makes it ineffective for discrimination and pattern recognition for “relaxation” despite its relationship to happiness. Another numerical feature only important in some contexts of change is “sunshine,” which was selected using *Original, First,* and *Pairwise*, but not *Previous*. Sunshine has a linear relationship to happiness over time which can sometimes be less impactful upon calculating Δs using differences between adjacent timepoints only.

EXPLANA can be used with cross-sectional data, which does not include Δs. To illustrate this, EXPLANA was applied to *SimFeatures* at timepoint 1. Perfect precision and recall were obtained, with six features selected: high and low salary, fish and dog as pets, healthy and unhealthy individuals (Supplemental Figure 3).

EXPLANA performance was then compared to an existing feature selection tool for LMS, called QIIME 2^38^ longitudinal^16^ feature-volatility. Because feature-volatility does not create Δ datasets (although other tools in q2-longitudinal can create Δ datasets) and does not automatically include categorical variables like EXPLANA, *SimMicrobiome* (the simulated microbiome, with no other study input variables) was analyzed using feature-volatility and EXPLANA (Table 1; Supplemental Figure 2). Default analyses for feature-volatility is on *Original*, and for EXPLANA on *Original, First, Previous* and *Pairwise*. All performance measures were substantially better for EXPLANA compared to feature-volatility, except for recall, which was 0.92 for EXPLANA *Original* and 1.00 for feature-volatility (Table 1). Of the 25 predictive microbes, 23 were selected by EXPLANA and 25 by feature-volatility. Of 175 not-predictive microbes, EXPLANA selected 5 FPs, while feature-volatility selected 155 FPs.

### Feature selection using a published study

EXPLANA was next applied to identify bacteria related to month-of-life in babies from the Early Childhood Antibiotics and Microbiome (ECAM) study^39^ which was also used to compare results to QIIME 2 longitudinal feature-volatility feature selection tool.^16,38^ The ECAM study generated 16S rRNA targeted sequencing data from monthly fecal samples collected from 43 babies over their first 2 years. Of 455 genera, 61 were selected by feature-volatility *Original*, and for EXPLANA, 37 for *Original,* 46 for *First*, 26 for *Previous*, and 96 for *Pairwise* (Figure 5). Of the 61 genera identified with feature-volatility, 25 were rejected by EXPLANA with *Original*. In total, there were 37 genera unique to EXPLANA analyses, with one identified using *Original*, which is an unidentified genus from the family *Methylobacteriaceae*. 36 unique genera were identified using *First, Previous* and *Pairwise* Δ datasets including *Paracoccus, Allobaculum, Anaerostipes,* and *Lactococcus,* which were each identified in two of the three Δ datasets.

**Figure 5.**
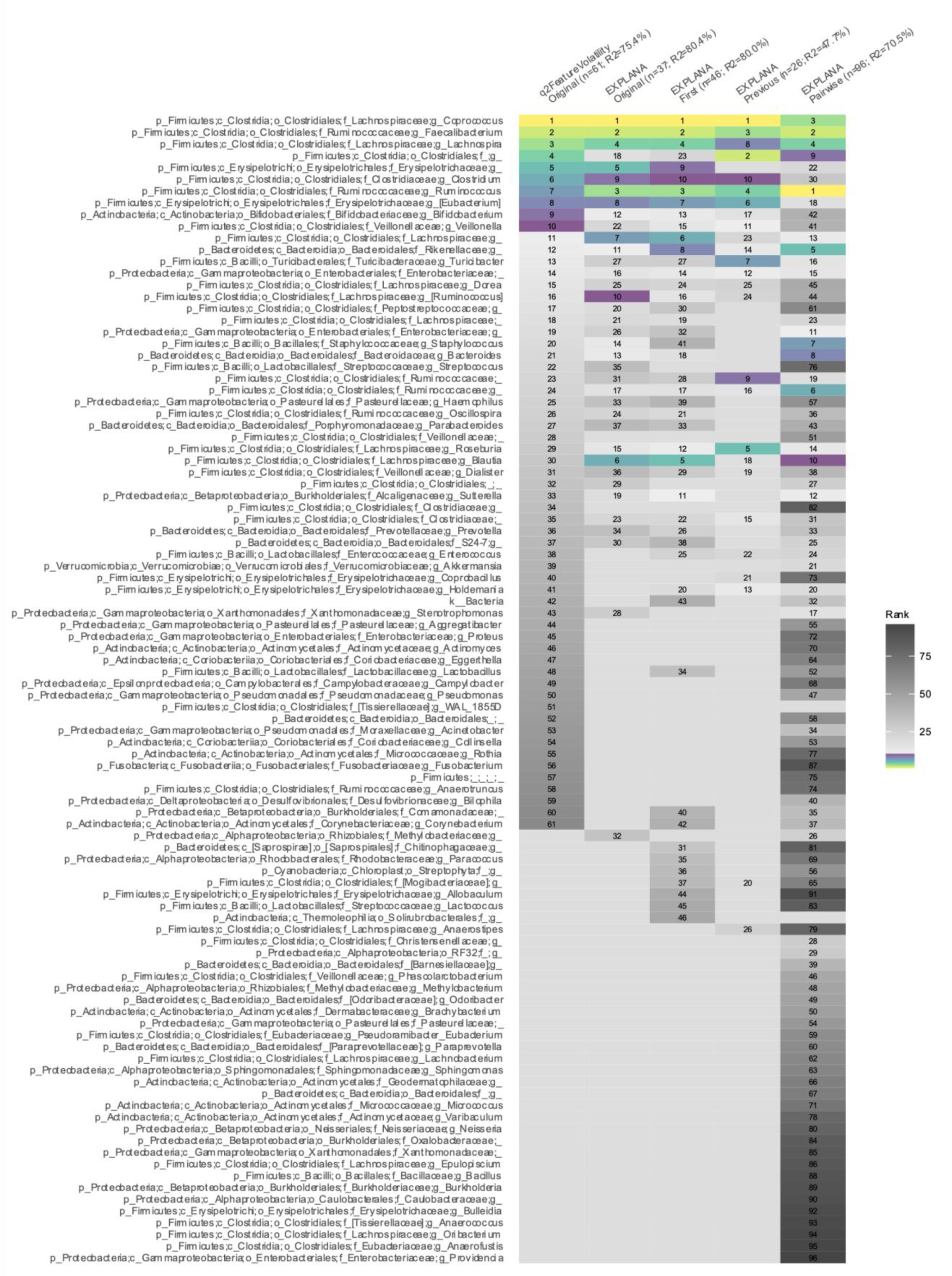
Bacterial genera related to month-of-life in newborns from the Early Childhood Antibiotics and Microbiome (ECAM) dataset selected using EXPLANA and QIIME 2 longitudinal feature-volatility.^16,38,39^. Features are ranked with one being the most important. Feature-volatility results are in first column (*Original* model), followed by EXPLANA for *Original, First, Previous* and *Pairwise*. Software tool, model and percent variation explained using R^2^ is shown on the X-axis. Both tools used 500 trees. EXPLANA had a feature fraction of 0.3, max depth of 7, 10 iterations of mixed-effects Random Forests (MERFs), and 100 BorutaSHAP trials (100% threshold; p=0.05). For feature-volatility, the feature fraction is a fixed parameter at 1.0 and uses Random Forest (RF) without mixed-effects. Top ten features per model are emphasized using a sequential multi-hue color palette from light to dark, and features after 10 are in grayscale from light to dark. *Paracoccus, Allobaculum, Anaerostipes* and *Lactococcus* genera were uniquely identified with EXPLANA using Δ datasets. *Blautia* had a much higher importance rank with EXPLANA compared to feature-volatility.

To demonstrate the versatility for EXPLANA to work with categorical variables using the ECAM dataset we identified categorical features related to month-of-life, while excluding the numerical microbiome data. Variables included delivery type (cesarean/vaginal), diet during first three months (breast/formula milk), sex (male/female), and antibiotic exposure (yes/no). Change in antibiotic use from no to yes (n_y) at later timepoints was positively related to month-of-life, indicating that babies are more likely to have an antibiotic treatment event as they age. (Supplemental Figure 4).

## Discussion

To address challenges with LMS analytics, we developed a feature selection tool for longitudinal data to expedite discovery. Supervised ML methods were implemented to identify features that relate to outcome values. For meaningful results, it was essential to use methods that not only identified features that were important for regression, but also implement tools such as Boruta, that evaluate whether they are more important than by chance, and SHAP, that provide evaluation of direction of change. These tools improve confidence that relevant features were selected, and facilitate hypothesis generation by explaining feature impact on dependent outcome variables.

EXPLANA had good performance with simulated data, as determined by selection or rejection of engineered features. The Δ datasets highlighted that change analyses can produce unique insights compared to *Original* longitudinal data without Δ calculations. Several studies have applied MERFs for feature selection,^40,41^ as used in EXPLANA, however, they did not use Δs, which could lead to a loss of valuable insights.

The four models (*Original, First, Previous* and *Pairwise*) can have strengths and limitations with feature selection. For example, *Previous* failed to select “sunshine” with *SimFeatures*, which had an engineered linear relationship to happiness over time, while the other three models selected it with a high importance rank. This finding is related to our observation that *Previous* had a low percent variation explained and recall for *SimMicrobiome* because the simulated predictive microbes and sunshine had a linear trend over time. Other temporal trends include quadratic, hockey stick, etc.^37^ *Previous* can miss predictive features when changes are minimized such as when the reference time is closer and predictor variables linearly relate to the outcome. This contrasts with *First*, where changes would be emphasized compared to baseline. A limitation of *Pairwise*, is that more comparisons are made and comparisons from overlapping time spans are not independent, so it is important to consider a higher chance of FPs, which we did observe. Therefore, stringent statistical parameters should be considered. For all simulations, *First* and *Original* had the best overall percent variation explained as expected for this study design consisting of engineered features with known changes from a baseline value.

Despite model limitations in particular instances, engineered features emphasized the importance in building models using *Original* and Δ datasets. Pill color “green_blue” was only able to be found in *Previous* and *Pairwise*, and demonstrated the ability for EXPLANA to use Δ datasets to find order-dependent categorical variables that impact an outcome (also seen with “no_yes” for antibiotics in the ECAM data), which are impossible to detect using *Original*. “Relaxation” is impossible to select in *First,* and was not selected, because it lacked variation compared to baseline values, it was selected by all other models. These features provide examples that demonstrate how different models, for different contexts of change, are needed to uncover time-dependent effects. The inclusion of ordered variables allows for modeling statistical interactions with time, which are often difficult to interpret, especially with non-linear approaches like Random Forests. By incorporating ordered variables in the automated change calculation step, and providing visualizations in the report, the workflow enhances interpretation of findings from longitudinal studies.

Dissimilarities between data included in each of the four models can create complications with interpreting feature selection results, such as dropping samples from Δ datasets due to missing timepoints. Another challenge with interpreting results from multiple models arises from including distance matrices, which can only be included in Δ datasets because they represent changes between samples. Thus, care should be taken with interpreting results obtained by different models within one report.

A total of 43 babies were included in the ECAM data reanalysis, which is a relatively small n. Other studies have demonstrated that RFs are useful for feature selection with sample sizes in the range of 25-35 samples. ^42–44^ The higher number of important genera selected using feature-volatility is likely due to a lack of statistical testing to reduce FPs as done with BorutaSHAP in EXPLANA. Indeed, many FPs were identified using feature-volatility with simulated microbiome data (*SimMicrobiome*) which contained a known amount of predictive and not predictive microbes. For the ECAM dataset, importance ranks differ for many genera between EXPLANA and feature-volatility leading to different conclusions about the degree of importance regarding developmental microbiome changes. Notably, the genera *Roseburia, Ruminococcus* and *Blautia* were ranked higher with EXPLANA compared to feature-volatility, and *Bifidobacterium* and *Veillonella* ranked lower using all four models (Figure 5). There were 37 unique genera found by EXPLANA, and not feature-volatility.

The tools first-differences and first-distances (for distance matrices) from q2-longitudinal^1^ can create Δs from continuous data, which can be used with feature-volatility. However, creation of Δs is not part of the feature-volatility feature-selection process, and comparing results from different models is cumbersome without comparative figures, such as the feature occurrences figure provided with EXPLANA (e.g., Figure 4a). Additionally, the incorporation of categorical Δs is a novel method not performed by other feature selection tools. The ML model used in feature-volatility is RF, which is not designed for repeated measures like MERFs used in EXPLANA. The importance score used in feature-volatility is Gini, which is biased when categorical and numerical variables are combined,^45^ while EXPLANA uses SHAP, which works well for this combination of feature types. Additionally, SHAP provides feature impact on the outcome, in addition to rank, as well as statistical testing from BorutaSHAP. Overall, analyzing the ECAM data with the mixed-effects models and statistical testing used in EXPLANA yielded a more meaningful set of bacterial genera and unique genera using Δ datasets.

There is no one-size-fits all model and it is challenging to understand which parameter adjustments will lead to optimal results. However, tuning of the algorithm can address some issues, especially the number of features available per decision tree split (a parameter that cannot be modified in feature-volatility), which is affected by the proportion of meaningful and collinear input variables. This workflow simplifies the process of comparing and testing different models using different arguments to identify which is the most effective model. Interpretation of exploratory analysis should be done with care, and *post-hoc* testing should be considered.

The barrier to performing data-driven feature selection for cross-sectional and LMS has been lessened by EXPLANA. Different applications are possible including focusing analysis on specific timepoints or segments of time within a longer study period, such as during plateau or active time periods. Additionally, stratifying by factors such as sex, geography, disease symptom, or a combination of factors could be worthwhile. EXPLANA also provides the opportunity to investigate different variables as inputs, sets of inputs, or as outcomes from prior hypotheses or from results of another exploratory analysis.

Overall, EXPLANA addresses many challenges of high-dimensional exploratory data analysis by combining existing tools and novel methods, and facilitates data-driven hypothesis generation.

## Methods

### Workflow overview

EXPLANA was developed using Snakemake^46^ to facilitate piping inputs and outputs from scripts written in different software languages, primarily R and Python (Figure 1). The workflow is executed from user-input arguments from a configuration file which pipes files to different scripts concluding with an .html report. The configuration file includes a list of datasets (microbiome, surveys, etc.) in long format (rows are samples; columns are features). First, individual datasets can be preprocessed through filters, dimensionality reduction, or transformation. If multiple files exist, they are merged to create the *Original* dataset. For longitudinal data, Δ datasets are computed (Figure 2). Finally, a feature selection algorithm is implemented by building a model from each of the four datasets (*Original, First Previous* and *Pairwise*): First, RFs^13^ or MERFs^15^ (for multiple samples per subject), are trained; Next, BorutaSHAP^36^ is used to rank features by importance if they perform better than expected by random chance, and determine feature impact on response. The final report includes figures, tables, and a written analytic summary.

Analyses were completed locally to ensure reasonable compute time for typical academic microbiome studies or those without server access. For 1000 features, 5 timepoints and 100 individuals, run time is less than 30 minutes using a MacBook Pro (Memory: 32 GB 2400 MHz DDR4; Processor: 2.9 GHz; 6-Core Intel Core i9).

### Software and data availability

Software is free for academic use. Workflow implementation and documentation can be found at www.explana.io and software, sample datasets and licensing at https://github.com/JTFouquier/explana.

### Configuration file

Each configuration file is associated with one analysis. Users modify a configuration file that specifies datasets, a response/outcome variable, sample identifier column, timepoint column, distance matrices (if applicable), optional dimensionality reduction steps prior to feature selection, and ML method decisions/arguments. Feature values as well as feature columns can be kept or dropped for individual datasets or for the merged *Original* dataset using small scripts within the configuration file.

### Preprocessing datasets

For each feature selection analysis, one or more dataset files can be used as needed. Each dataset can be preprocessed, which may include dropping features or feature values, on a per dataset basis. Dimensionality reduction can be performed prior to feature selection using principal components analysis (PCA), transformation or filters. PCA is used on a set of related variables to capture the maximum variance using fewer variables. Short scripts can be added to the configuration file to modify each dataset or the complete dataset after merging individual datasets.

### Dataset integration

After preprocessing, individual datasets are merged using sample identifier column to create the “*Original*” dataset. The *Original* dataset is named accordingly because it contains original values of features that may have been sampled over time (i.e., *Original* does not include intra-individual changes/differences between timepoints like the Δ datasets). Data integration prioritizes samples in the top/first dataset. This means additional samples in other datasets will not be included. For some analyses, merging data prior to implementation may be simpler.

### Delta (Δ) dataset creation

For longitudinal analyses, the *Original* dataset is used to compute three Δ datasets, *First*, *Previous*, and *Pairwise* by calculating feature changes over time, per subject, using different reference points (Figure 2). Δ dataset calculations: For *First*, compared to baseline/first; for *Previous*, compared to previous timepoint; for *Pairwise*, all pairwise comparisons between timepoints. For two timepoint studies, only *Original* and *First* are needed.

For categorical variables, the order of categorical values for each subject at both timepoints per comparison is tracked (e.g., for pill color, if T2 is green and T3 is blue, T2_T3 is green_blue). For numerical variables, reference values are subtracted from the later timepoint (e.g., if T2 is 3 and T3 is 7, Δ = 4).

Timepoint is numerical for *Original* to provide information about order of events, and categorical for Δ datasets due to overlap in timepoints (i.e., T1_T2 and T1_T3 overlap each other at T1_T2). This overlap can be thought of as though time were categorical rather than an abstract concept. In other words, if T1, T2 and T3 were recoded as A, B, and C, respectively, the comparisons A_B, A_C and B_C are potentially interesting.

### Feature selection algorithm

Feature selection is performed using all four models, as needed. For categorical features, unique values/classes per feature are encoded to binary features (labeled as “ENC”), where feature presence in a sample is 1 and absence is 0. This enables selection of uncommon feature values that influence an outcome variable.

Next, RF regression is performed to select features related to the outcome. When more than one measurement per subject exists, MERF is used. Both use Scikit-Learn^47^ RandomForestRegressor as the fixed effects forest. Boruta^34^ is a method that uses shuffled versions of input features to assess whether importance scores are better than random chance. Features are categorized as accepted, tentative or rejected. BorutaSHAP^36^ is implemented because it works with the unique properties of SHAP (SHapley Additive exPlanations),^35^ which provides feature ranks and estimated impact on the outcome.

Rejected features are dropped, RF (or MERF) is re-run, and visualizations are generated without irrelevant features that might hide true signal from important features. Final percent variation explained comes from OOB out-of-bag (OOB) scores from the complete-feature forest. For additional context, the percent variation explained by the reduced-feature forest is also provided. OOB scores are used for internal validation and created from leaving out some samples for each decision tree in the forest during training and comparing the tree’s results to the real outcome values for the samples left out.

### Report Details

The result of each analysis is an interactive .html report that includes figures, tables, links to directories for data exploration, links to PubMed for researching findings, and written descriptions of the analytic process (Figure 3). A methods section is dynamically created based on user inputs or defaults, which can be included in manuscripts. A feature occurrence figure summarizes feature ranks for all models, followed by model-specific sections and impact on outcome is provided for categorical features because positive instances per sample are clear, while numerical feature relationships are more complex (e.g., a hockey curve pattern). Figures include SHAP plots that explain feature impact on outcome.

### Simulation study design

A simulation study on happiness was designed to facilitate performance testing using variables that are either predictive or not predictive of happiness. Variables and their effects on the outcome are in Table 1. All individuals started with the same happiness score and effects from all features were used to update each subject’s happiness score at each timepoint. Predictive features had effect values stored in a different column labeled with the suffix “_effect”. For example, the “salary” column contained values “high” or “low” and had a corresponding “salary_effect” column with numerical values reflecting effects on happiness. All effect column values were added to original happiness scores and columns labeled “effect” were removed before feature selection. This way, engineered effects were contained in the happiness value, and predictive features can be identified if they corresponded to the effect.

R package faux^48^ was used to add subject random effects, five timepoints, and a control and test group. The test group was simulated to linearly increase with a positive slope of 30 (correlation coefficient of 0.7 and SD = 5) to simulate a treatment effect that improved happiness.

To create longitudinal microbiome simulations containing differentially abundant and not differentially abundant microbes, microbiomeDASim^37^ was used with a first-order autoregressive correlation structure that linearly increased with slope 30 to correlate with happiness (Correlation coefficient = 0.7; standard deviation = 5).

Eight data distributions were used with random, not predictive variables. Normal, Bernoulli, Binomial, Poisson, Exponential, Gamma, Weibull^49^ and Dirichlet. The number of random variables is indicated in the dataset name. Accordingly, for *SimFeaturesMicrobiome0*, *SimFeaturesMicrobiome500* and *SimFeaturesMicrobiome1000*, the number of random variables is 0, 500, and 1000, respectively.

When using EXPLANA, arguments were set based on recommendations of the underlying tools or from previous studies on hyperparameter tuning,^50,51^ which included for MERF, 300 trees, 0.3 feature fraction for decision tree splits with a max depth of 7 and 10 MERF iterations and 100 BorutaSHAP trials were run (100% importance threshold; *p* = 0.05).

### Performance evaluation

Performance was assessed using simulation studies and F1-scores and AUC metrics (Supplemental Figure 1). Recall (TP rate): proportion of predictive features correctly selected (Recall = TP/(TP+FN)). Precision: proportion of all selected features that are truly predictive (Precision = TP/(TP+FP) An F1-score is calculated using precision and recall 2*(Precision * Recall) / (Precision + Recall), as well as AUC, which plots the TP rate against the FP rate.

### Early Childhood Antibiotics and Microbiome (ECAM) dataset analysis

To explore feature selection results using published data, 455 genera from the ECAM dataset were used because it was also used to demonstrate QIIME2^38^ longitudinal feature-volatility^16^ functionality. Metadata was filtered to remove duplicate months to facilitate Δ calculations performed by EXPLANA. 500 trees were used for both tools. EXPLANA arguments: Feature fraction of 0.3, max depth = 7, with 10 MERF iterations and 100 BorutaSHAP trials (100% threshold; p = 0.05). Q2 feature-volatility arguments: Feature fraction of 1.0 (cannot modify) and 5 k-fold cross-validations.

## Acknowledgements

We would like to acknowledge the following individuals from The University of Colorado, Anschutz Medical Campus: Laura Saba, Michael Strong, Katerina Kechris, Emily Mastej, Nichole Nusbacher, Casey Martin, Mallory Karr, Brook Santangelo, John Sterrett, Angela Sofia Burkhart Colorado, Erik Serrano, Elena Wall, Madison Apgar, and Luis Treviso. We would also like to thank Scott Kelley (San Diego State University), and Alba Talavera-Rodríguez and Sergio Serrano-Villar (University Hospital Ramón y Cajal, Spain) for feedback and suggestions.

## Supplemental Figures

**Supplemental Figure 1.**
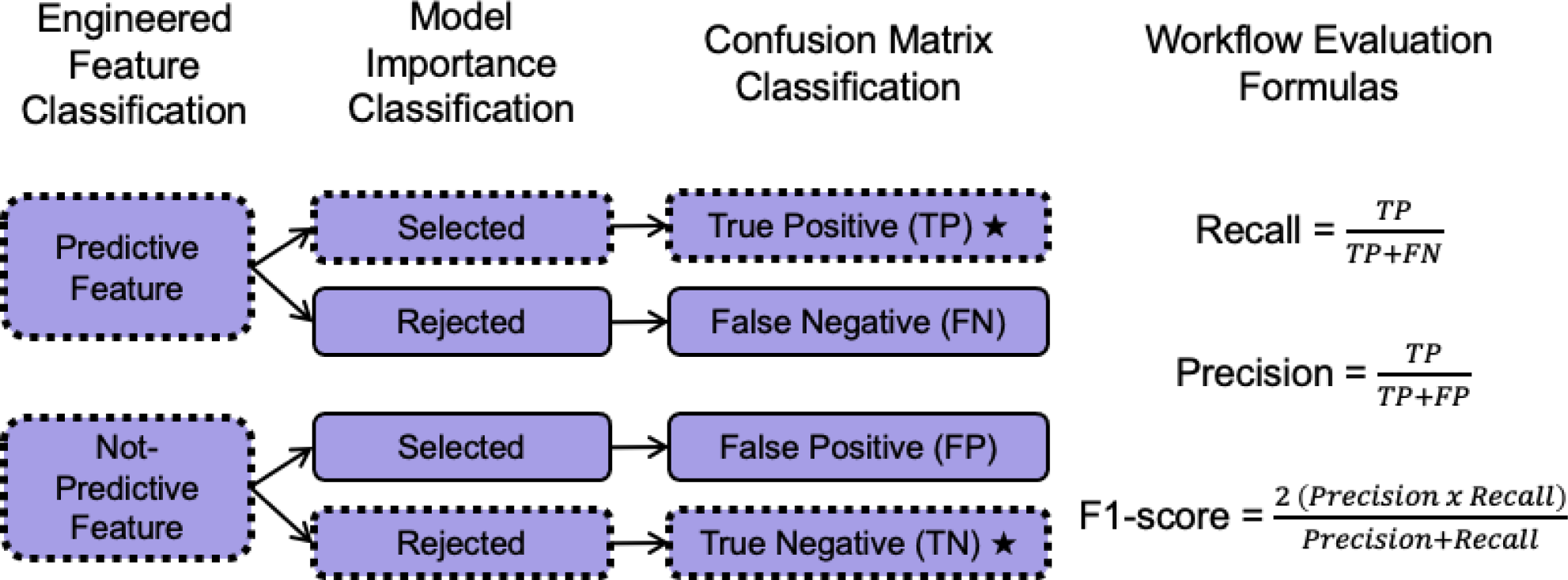
Classification diagram for engineered predictive and not predictive features used for model performance evaluation. The simulation study contains features with a relationship to the outcome variable (predictive features) and without (not-predictive features). Dashed lines and stars indicate the correct classification paths for engineered features. Recall (TP rate) is the proportion of predictive features correctly selected (Recall = TP/(TP+FN)). Precision is the proportion of all selected features that are truly predictive (Precision = TP/(TP+FP)). An F1-score is calculated using precision and recall (2*(Precision * Recall) / (Precision + Recall)). TP = true positive, FP = false positive.

**Supplemental Figure 2.**
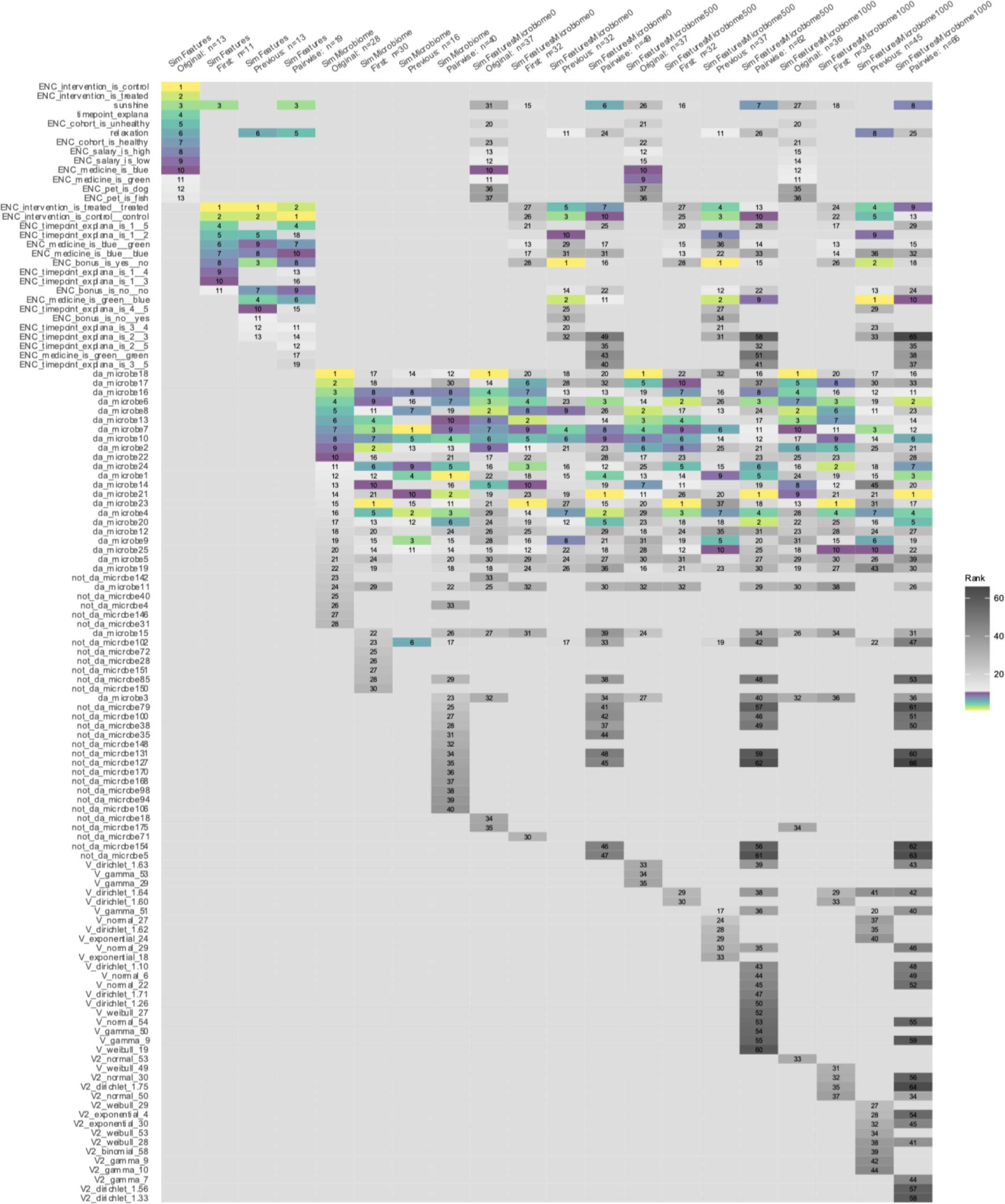
Features related to happiness selected using EXPLANA with five variations of simulated longitudinal microbiome datasets. The *SimFeatures* dataset is a simulated longitudinal intervention with 100 individuals sampled over five timepoints. *SimMicrobiome* is a simulated longitudinal microbiome dataset created using MicrobiomeDASim^37^ with a ratio of 25 differentially abundant microbes to 175 not differentially abundant. See Methods and Supplemental Table 1 for detailed description of the simulation study design. *SimFeaturesMicrobiome0*, *SimFeaturesMicrobiome500*, and *SimFeaturesMicrobiome1000* are dataset variations that include a simulated microbiome, study variables and random variables from a variety of data distributions with no relationship to the outcome. The number in the dataset name represents the number of random variables included. 300 trees were used, with a feature fraction of 0.3, max depth of 7, with 10 iterations of mixed-effects Random Forests (MERFs), and 100 BorutaSHAP trials (100% importance threshold at p=0.05). Top ten features per model are emphasized using a sequential multi-hue color palette from light to dark, and features after 10 are in grayscale from light to dark. Notable features include “sunshine,” which was selected in *Original, First* and *Pairwise*, but not selected in *Previous*; “relaxation” which was not selected in *First* but was selected in *Original, Previous* and *Pairwise*; and “green_blue,” an order-dependent categorical feature that impacted the response and is only able to be found using delta Δ datasets.

**Supplemental Figure 3.**
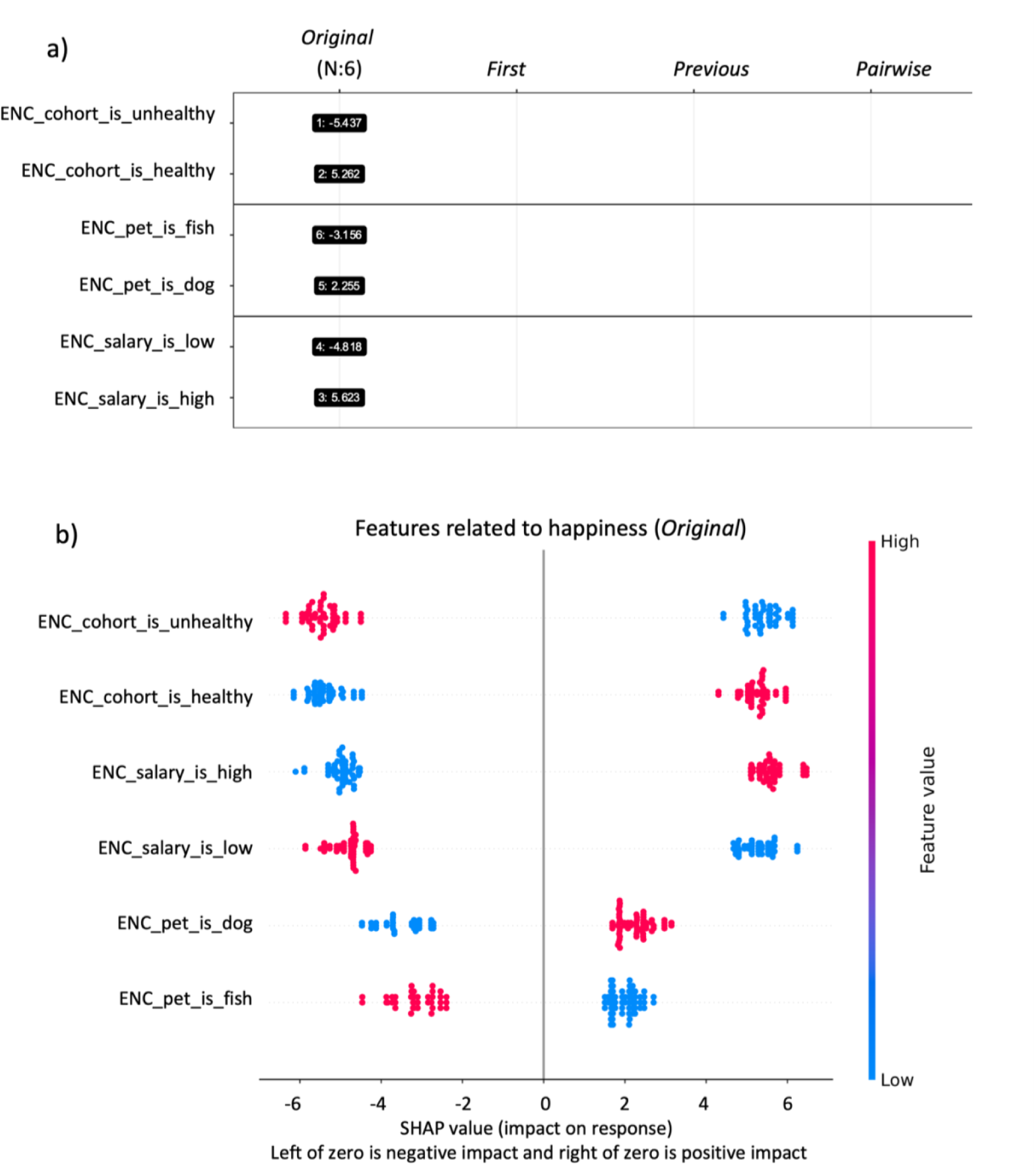
Cross-sectional feature-selection results using EXPLANA with baseline values only from simulated happiness dataset. *SimFeatures* dataset was used at timepoint 1 and cohort, salary and pet were selected as important. (a) Feature occurrence diagram displaying rank and SHAP value. Unhealthy individuals are ranked 1 and have a −5.4 impact on “happiness” and healthy individuals are ranked 2 and have a 5.2 impact. (b) SHAP summary beeswarm plot where each point represents one sample, and the horizontal position indicates impact on the outcome as indicated on the x-axis. Points to the left indicate a negative impact, and points to the right indicate a positive impact. The colors represent the selected feature values, where red is larger, and blue is smaller. For binary encoded features (‘ENC’) red is yes/1 and blue is no/0. Features are ordered by largest to smallest impact on the response. As shown, low salary negatively impacts happiness and having a dog positively impacts happiness.

**Supplemental Figure 4.**
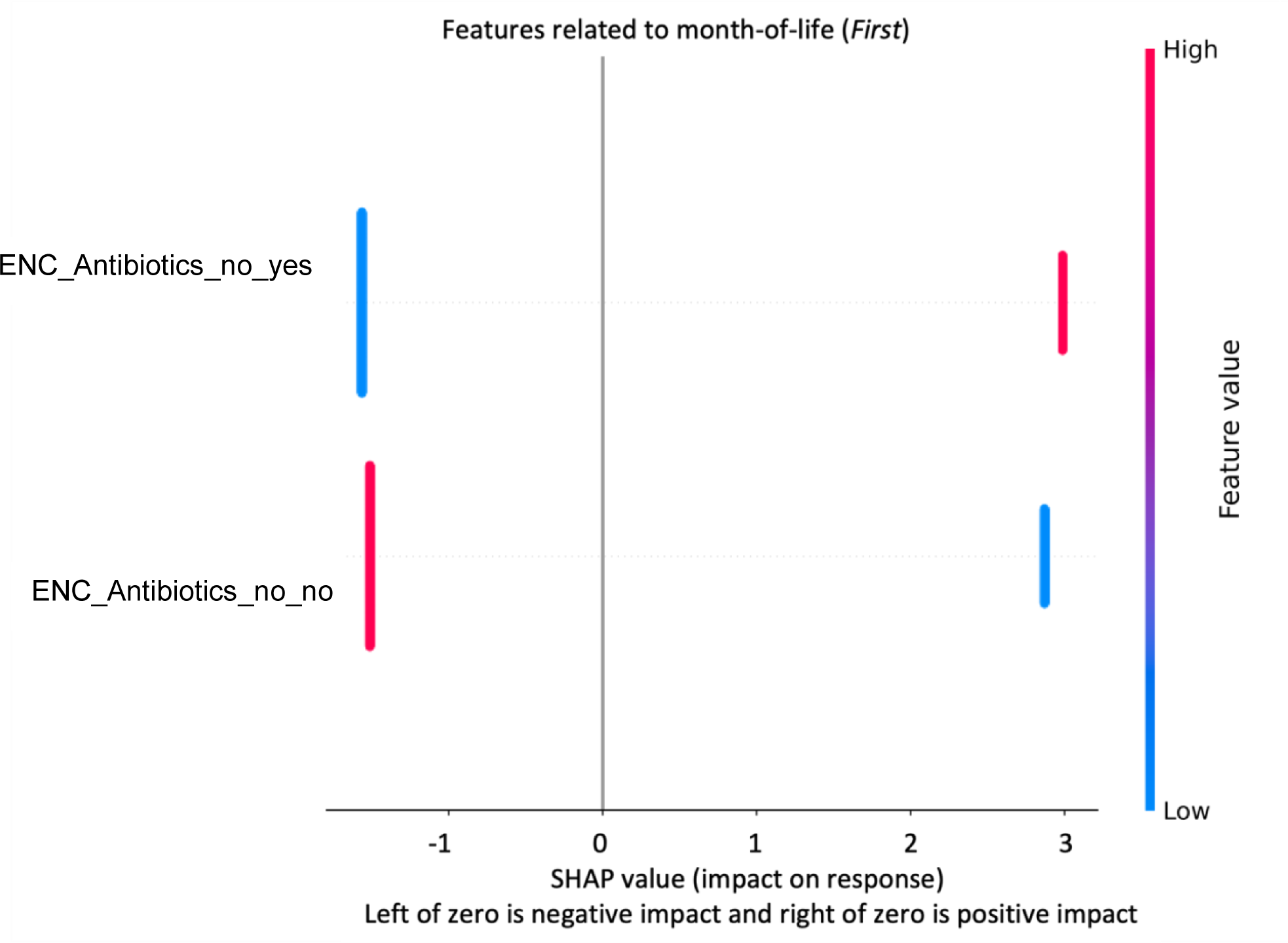
Heatmap of bacterial genera predictive of month-of-life in newborns selected by EXPLANA using only categorical features from the Early Childhood and Microbiome (ECAM) dataset. RF was used with 500 trees, a feature fraction of 0.2, max depth of 7, and 100 BorutaSHAP trials (100% threshold; p=0.05). Each point represents one sample, and the horizontal position indicates impact on the outcome as indicated on the x-axis. Points to the left indicate a negative impact, and points to the right indicate a positive impact. The colors represent the selected feature values, where red is larger, and blue is smaller. For binary encoded features (‘ENC’) red is yes/1 and blue is no/0.

## Supplemental Tables

**Supplemental Table 1.**
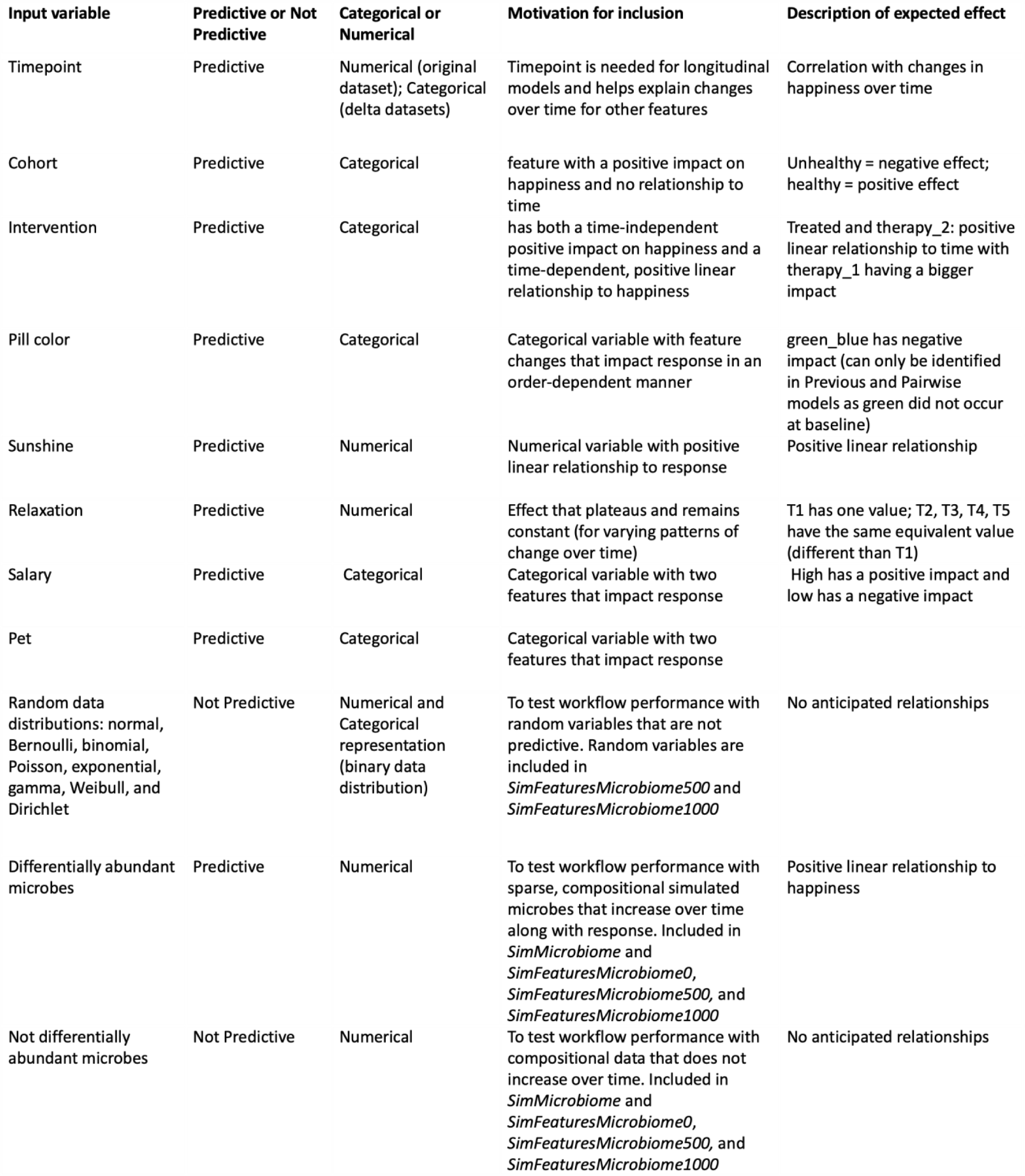
Engineered predictive and not-predictive input features used in happiness simulation studies.

## References

1. Santiago-Rodriguez, T. M. & Hollister, E. B. Multi ‘omic data integration: A review of concepts, considerations, and approaches. Seminars in Perinatology 45, 151456 (2021).

2. Ursell, L. K., Metcalf, J. L., Parfrey, L. W. & Knight, R. Defining the human microbiome. Nutrition Reviews 70, S38–S44 (2012).

3. Hrdlickova, R., Toloue, M. & Tian, B. RNA-Seq methods for transcriptome analysis. WIREs RNA 8, e1364 (2017).

4. Zamboni, N., Saghatelian, A. & Patti, G. J. Defining the Metabolome: Size, Flux, and Regulation. Molecular Cell 58, 699–706 (2015).

5. Maruvada, P., Leone, V., Kaplan, L. M. & Chang, E. B. The Human Microbiome and Obesity: Moving beyond Associations. Cell Host & Microbe 22, 589–599 (2017).

6. Valles-Colomer, M. et al. The neuroactive potential of the human gut microbiota in quality of life and depression. Nature microbiology 4, 623–632 (2019).

7. Krajmalnik-Brown, R., Lozupone, C., Kang, D.-W. & Adams, J. B. Gut bacteria in children with autism spectrum disorders: challenges and promise of studying how a complex community influences a complex disease. Microbial Ecology in Health and Disease 26, 26914 (2015).

8. Rebersek, M. Gut microbiome and its role in colorectal cancer. BMC cancer 21, 1–13 (2021).

9. Zhuang, H. et al. Dysbiosis of the gut microbiome in lung cancer. Frontiers in Cellular and Infection Microbiology 9, 112 (2019).

10. Williams, B., Landay, A. & Presti, R. M. Microbiome alterations in HIV infection a review. Cellular microbiology 18, 645–651 (2016).

11. Witkowski, M., Weeks, T. L. & Hazen, S. L. Gut microbiota and cardiovascular disease. Circulation research 127, 553–570 (2020).

12. Linear Mixed-Effects Models: Basic Concepts and Examples. in Mixed-Effects Models in S and S-PLUS (eds. Pinheiro, J. C. & Bates, D. M.) 3–56 (Springer, New York, NY, 2000). doi:10.1007/0-387-22747-4_1.

13. Breiman, L. Random Forests -- Random Features.

14. Díaz-Uriarte, R. & Alvarez de Andrés, S. Gene selection and classification of microarray data using random forest. BMC Bioinformatics 7, 3 (2006).

15. Hajjem, A., Bellavance, F. & Larocque, D. Mixed-effects random forest for clustered data. Journal of Statistical Computation and Simulation 84, 1313–1328 (2014).

16. Bokulich, N. A., et al. q2-longitudinal: Longitudinal and Paired-Sample Analyses of Microbiome Data. mSystems 3, 10.1128/msystems.00219-18 (2018).

17. Frey, D. L. et al. Changes in Microbiome Dominance Are Associated With Declining Lung Function and Fluctuating Inflammation in People With Cystic Fibrosis. Front. Microbiol. 13, (2022).

18. Ferrocino, I. et al. Changes in the gut microbiota composition during pregnancy in patients with gestational diabetes mellitus (GDM). Sci Rep 8, 12216 (2018).

19. Meslier, V. et al. Mediterranean diet intervention in overweight and obese subjects lowers plasma cholesterol and causes changes in the gut microbiome and metabolome independently of energy intake. Gut 69, 1258–1268 (2020).

20. Rodenes-Gavidia, A. et al. An insight into the functional alterations in the gut microbiome of healthy adults in response to a multi-strain probiotic intake: a single arm open label trial. Front Cell Infect Microbiol 13, 1240267 (2023).

21. Zhang, L., Luo, H. & Kang, G. Longitudinal study of physical activity with various methods in maintenance hemodialysis patients. Hemodialysis International 25, 249–256 (2021).

22. Fouquier, J. et al. The Gut Microbiome in Autism: Study-Site Effects and Longitudinal Analysis of Behavior Change. mSystems 6, e00848–20 (2021).

23. Twisk, J. W. R. *Applied Longitudinal Data Analysis for Epidemiology: A Practical Guide*. (Cambridge University Press, Cambridge, 2013). doi:10.1017/CBO9781139342834.

24. Tartini, R., Steinbrunn, W., Kappenberger, L. & Meyer, U. A. Dangerous interaction between amiodarone and quinidine. The Lancet 319, 1327–1329 (1982).

25. Vassallo, P. & Trohman, R. G. Prescribing Amiodarone: An Evidence-Based Review of Clinical Indications. JAMA 298, 1312–1322 (2007).

26. Mullen, S. A. et al. Precision therapy for epilepsy due to KCNT1 mutations: A randomized trial of oral quinidine. Neurology 90, e67–e72 (2018).

27. Queen, O. & Emrich, S. J. LASSO-based feature selection for improved microbial and microbiome classification. in 2021 IEEE International Conference on Bioinformatics and Biomedicine (BIBM) 2301–2308 (2021). doi:10.1109/BIBM52615.2021.9669485.

28. Wang, C., Hu, J., Blaser, M. J. & Li, H. Microbial trend analysis for common dynamic trend, group comparison, and classification in longitudinal microbiome study. BMC Genomics 22, 667 (2021).

29. Calle, M. L., Pujolassos, M. & Susin, A. coda4microbiome: compositional data analysis for microbiome cross-sectional and longitudinal studies. BMC Bioinformatics 24, 82 (2023).

30. Lee, K. H., Coull, B. A., Moscicki, A.-B., Paster, B. J. & Starr, J. R. Bayesian variable selection for multivariate zero-inflated models: Application to microbiome count data. Biostatistics 21, 499–517 (2018).

31. Bodein, A., Scott-Boyer, M.-P., Perin, O., Lê Cao, K.-A. & Droit, A. timeOmics: an R package for longitudinal multi-omics data integration. Bioinformatics 38, 577–579 (2022).

32. Gloor, G. B., Macklaim, J. M., Pawlowsky-Glahn, V. & Egozcue, J. J. Microbiome Datasets Are Compositional: And This Is Not Optional. Frontiers in Microbiology 8, (2017).

33. Lozupone, C., Lladser, M. E., Knights, D., Stombaugh, J. & Knight, R. UniFrac: an effective distance metric for microbial community comparison. ISME J 5, 169–172 (2011).

34. Kursa, M. B. & Rudnicki, W. R. Feature Selection with the Boruta Package. Journal of Statistical Software 36, 1–13 (2010).

35. Lundberg, S. M. & Lee, S.-I. A Unified Approach to Interpreting Model Predictions. In Advances in Neural Information Processing Systems vol. 30 (Curran Associates, Inc., 2017).

36. Keany, E. BorutaSHAP. (2021).

37. Williams, J., Bravo, H. C., Tom, J. & Paulson, J. N. microbiomeDASim: Simulating longitudinal differential abundance for microbiome data. F1000Res 8, 1769 (2020).

38. Bolyen, E. et al. Reproducible, interactive, scalable and extensible microbiome data science using QIIME 2. Nat Biotechnol 37, 852–857 (2019).

39. Bokulich, N. A. et al. Antibiotics, birth mode, and diet shape microbiome maturation during early life. Science Translational Medicine 8, 343ra82–343ra82 (2016).

40. Zeamer, A. L. et al. Association between microbiome and the development of adverse posttraumatic neuropsychiatric sequelae after traumatic stress exposure. Transl Psychiatry 13, 1–14 (2023).

41. Haran, J. P. et al. The high prevalence of Clostridioides difficile among nursing home elders associates with a dysbiotic microbiome. Gut Microbes 13, 1897209 (2021).

42. Hassan, S. S., Farhan, M., Mangayil, R., Huttunen, H. & Aho, T. Bioprocess data mining using regularized regression and random forests. BMC Syst Biol 7, S5 (2013).

43. Ward, D., Miller, R. & Nikolaev, A. Evaluating three stuttering assessments through network analysis, random forests and cluster analysis. Journal of Fluency Disorders 67, 105823 (2021).

44. Luan, J., Zhang, C., Xu, B., Xue, Y. & Ren, Y. The predictive performances of random forest models with limited sample size and different species traits. Fisheries Research 227, 105534 (2020).

45. Strobl, C., Boulesteix, A.-L., Zeileis, A. & Hothorn, T. Bias in random forest variable importance measures: Illustrations, sources and a solution. BMC Bioinformatics 8, 25 (2007).

46. Snakemake—a scalable bioinformatics workflow engine | Bioinformatics | Oxford Academic. https://academic.oup.com/bioinformatics/article/28/19/2520/290322.

47. Pedregosa, F. et al. Scikit-learn: Machine Learning in Python. Journal of Machine Learning Research 12, 2825–2830 (2011).

48. DeBruine, L. faux: Simulation for Factorial Designs. Zenodo 10.5281/ZENODO.2669586 (2023).

49. K, J. 7 Statistical Distributions that every Data Scientist should know— with intuitive explanations. Medium https://towardsdatascience.com/7-statistical-distributions-that-every-data-scientist-should-know-with-intuitive-explanations-bf967db81f0b (2020).

50. Weerts, H. J. P., Mueller, A. C. & Vanschoren, J. Importance of Tuning Hyperparameters of Machine Learning Algorithms. Preprint at http://arxiv.org/abs/2007.07588 (2020).

51. Probst, P., Wright, M. N. & Boulesteix, A.-L. Hyperparameters and tuning strategies for random forest. WIREs Data Mining and Knowledge Discovery 9, e1301 (2019).

