## Supplemental for "EXPLANA: A user-friendly workflow for EXPLoratory ANAlysis and feature selection in cross-sectional and longitudinal microbiome studies"

### Supplemental Figures

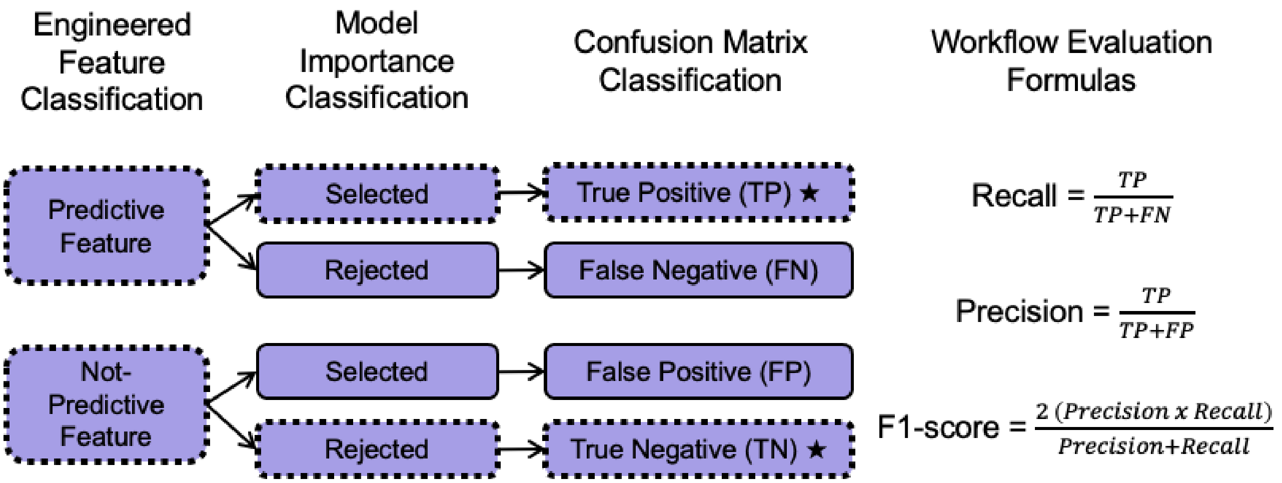


**Supplemental Figure 1. Classification diagram for engineered predictive and not predictive features used for model performance evaluation.** The simulation study contains features with a relationship to the outcome variable (predictive features) and without (not-predictive features). Dashed lines and stars indicate the correct classification paths for engineered features. Recall (TP rate) is the proportion of predictive features correctly selected (Recall = TP/(TP+FN)). Precision is the proportion of all selected features that are truly predictive (Precision = TP/(TP+FP)). An F1-score is calculated using precision and recall (2*(Precision * Recall) / (Precision + Recall)). TP = true positive, FP = false positive.

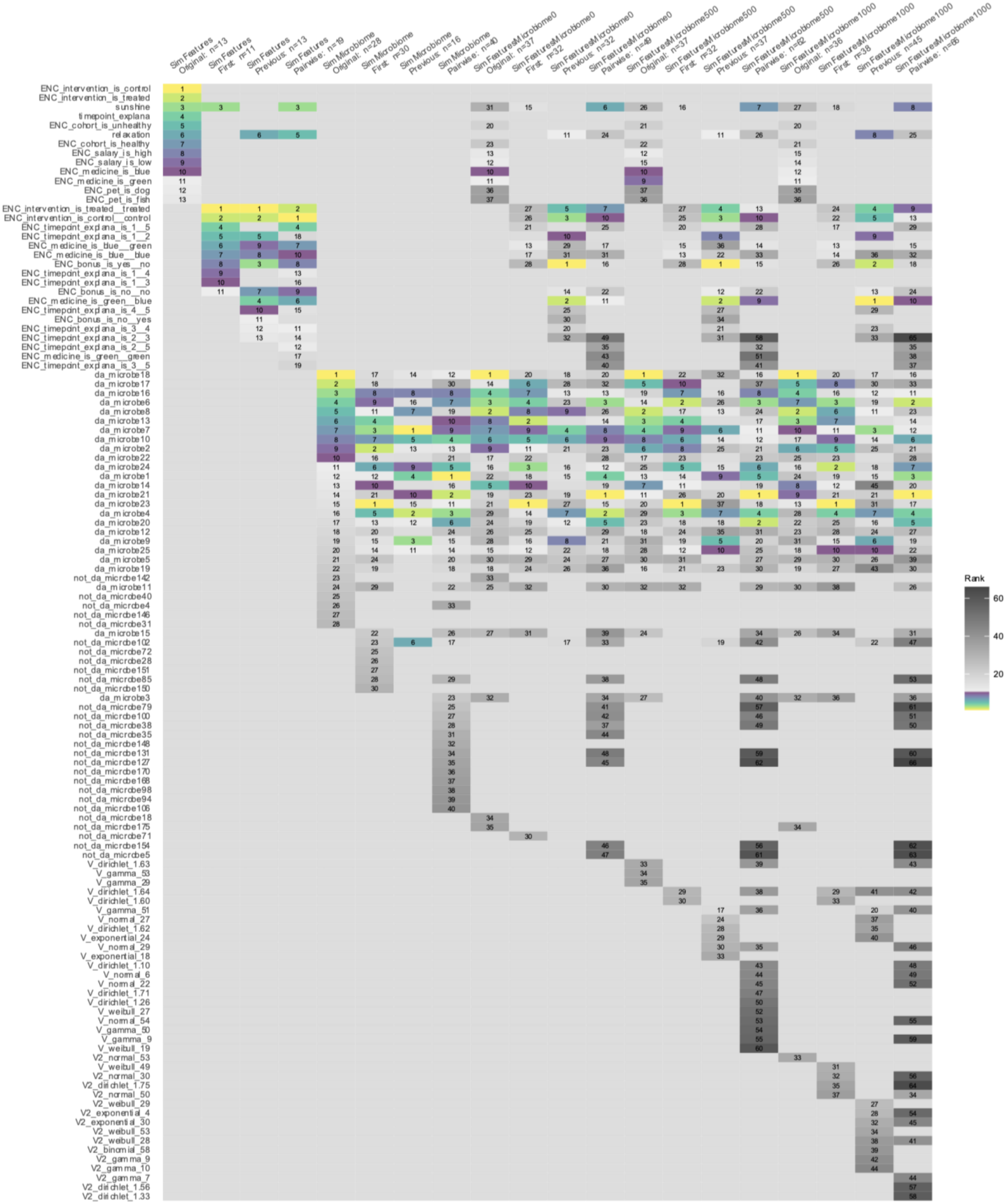

Sim Features Original: n=13
Sim Features First: n=11
Sim Features Previous: n=13
Sim Features Pairwise: n=19
Sim Microbiome Original: n=28
Sim Microbiome First: n=30
Sim Microbiome Previous: n=16
Sim Microbiome Pairwise: n=40
SimFeaturesMicrobiome0 Original: n=37
SimFeaturesMicrobiome0 First: n=32
SimFeaturesMicrobiome0 Previous: n=32
SimFeaturesMicrobiome0 Pairwise: n=49
SimFeaturesMicrobiome500 Original: n=37
SimFeaturesMicrobiome500 First: n=32
SimFeaturesMicrobiome500 Previous: n=37
SimFeaturesMicrobiome500 Pairwise: n=62
SimFeaturesMicrobiome1000 Original: n=36
SimFeaturesMicrobiome1000 First: n=38
SimFeaturesMicrobiome1000 Previous: n=45
SimFeaturesMicrobiome1000 Pairwise: n=66
ENC_intervention_is_control
ENC_intervention_is_treated
sunshine
timepoint_explana
ENC_cohort_is_unhealthy
relaxation
ENC_cohort_is_healthy
ENC_salary_is_high
ENC_salary_is_low
ENC_medicine_is_blue
ENC_medicine_is_green
ENC_pet_is_dog
ENC_pet_is_fish
ENC_intervention_is_treated__treated
ENC_intervention_is_control__control
ENC_timepoint_explana_is_1__5
ENC_timepoint_explana_is_1__2
ENC_medicine_is_blue__green
ENC_medicine_is_blue__blue
ENC_bonus_is_yes__no
ENC_timepoint_explana_is_1__4
ENC_timepoint_explana_is_1__3
ENC_bonus_is_no__no
ENC_medicine_is_green__blue
ENC_timepoint_explana_is_4__5
ENC_bonus_is_no__yes
ENC_timepoint_explana_is_3__4
ENC_timepoint_explana_is_2__3
ENC_timepoint_explana_is_2__5
ENC_medicine_is_green__green
ENC_timepoint_explana_is_3__5
da_microbe18
da_microbe17
da_microbe16
da_microbe6
da_microbe8
da_microbe13
da_microbe7
da_microbe10
da_microbe2
da_microbe22
da_microbe24
da_microbe1
da_microbe14
da_microbe21
da_microbe23
da_microbe4
da_microbe20
da_microbe12
da_microbe9
da_microbe25
da_microbe5
da_microbe19
not_da_microbe142
da_microbe11
not_da_microbe40
not_da_microbe4
not_da_microbe146
not_da_microbe31
da_microbe15
not_da_microbe102
not_da_microbe72
not_da_microbe28
not_da_microbe151
not_da_microbe85
not_da_microbe150
da_microbe3
not_da_microbe79
not_da_microbe100
not_da_microbe38
not_da_microbe35
not_da_microbe148
not_da_microbe131
not_da_microbe127
not_da_microbe170
not_da_microbe168
not_da_microbe98
not_da_microbe94
not_da_microbe106
not_da_microbe18
not_da_microbe175
not_da_microbe71
not_da_microbe154
not_da_microbe5
V_dirichlet_1.63
V_gamma_53
V_gamma_29
V_dirichlet_1.64
V_dirichlet_1.60
V_gamma_51
V_normal_27
V_dirichlet_1.62
V_exponential_24
V_normal_29
V_exponential_18
V_dirichlet_1.10
V_normal_6
V_normal_22
V_dirichlet_1.71
V_dirichlet_1.26
V_weibull_27
V_normal_54
V_gamma_50
V_gamma_9
V_weibull_19
V2_normal_53
V_weibull_49
V2_normal_30
V2_dirichlet_1.75
V2_normal_50
V2_weibull_29
V2_exponential_4
V2_exponential_30
V2_weibull_53
V2_weibull_28
V2_binomial_58
V2_gamma_9
V2_gamma_10
V2_gamma_7
V2_dirichlet_1.56
V2_dirichlet_1.33
Rank
60
40
20

**Supplemental Figure 2. Features related to happiness selected using EXPLANA with five variations of simulated longitudinal microbiome datasets.** The *SimFeatures* dataset is a simulated longitudinal intervention with 100 individuals sampled over five timepoints. *SimMicrobiome* is a simulated longitudinal microbiome dataset created using MicrobiomeDASim[37] with a ratio of 25 differentially abundant microbes to 175 not differentially abundant. See Methods and Supplemental Table 1 for detailed description of the simulation study design. *SimFeaturesMicrobiome0*, *SimFeaturesMicrobiome500*, and *SimFeaturesMicrobiome1000* are dataset variations that include a simulated microbiome, study variables and random variables from a variety of data distributions with no relationship to the outcome. The number in the dataset name represents the number of random variables included. 300 trees were used, with a feature fraction of 0.3, max depth of 7, with 10 iterations of mixed-effects Random Forests (MERFs), and 100 BorutaSHAP trials (100% importance threshold at p=0.05). Top ten features per model are emphasized using a sequential multi-hue color palette from light to dark, and features after 10 are in grayscale from light to dark. Notable features include “sunshine,” which was selected in *Original, First* and *Pairwise*, but not selected in *Previous*; “relaxation” which was not selected in *First* but was selected in *Original, Previous* and *Pairwise*; and “green_blue,” an order-dependent categorical feature that impacted the response and is only able to be found using delta $\Delta$ datasets.

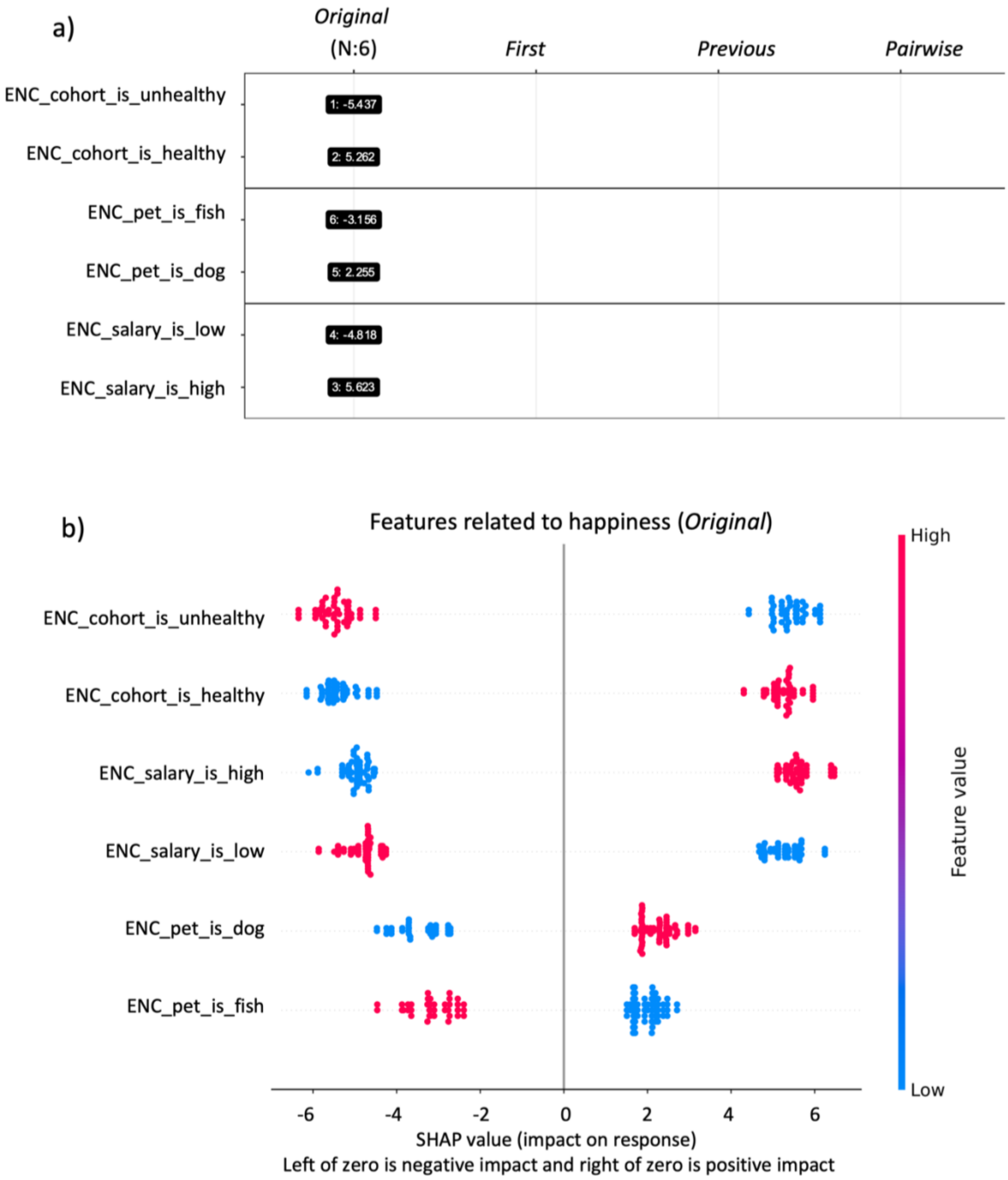


**Supplemental Figure 3. Cross-sectional feature-selection results using EXPLANA with baseline values only from simulated happiness dataset.** *SimFeatures* dataset was used at timepoint 1 and cohort, salary and pet were selected as important. (a) Feature occurrence diagram displaying rank and SHAP value. Unhealthy individuals are ranked 1 and have a -5.4 impact on “happiness” and healthy individuals are ranked 2 and have a 5.2 impact. (b) SHAP summary beeswarm plot where each point represents one sample, and the horizontal position indicates impact on the outcome as indicated on the x-axis. Points to the left indicate a negative impact, and points to the right indicate a positive impact. The colors represent the selected feature values, where red is larger, and blue is smaller. For binary encoded features (‘ENC’) red is yes/1 and blue is no/0. Features are ordered by largest to smallest impact on the response. As shown, low salary negatively impacts happiness and having a dog positively impacts happiness.

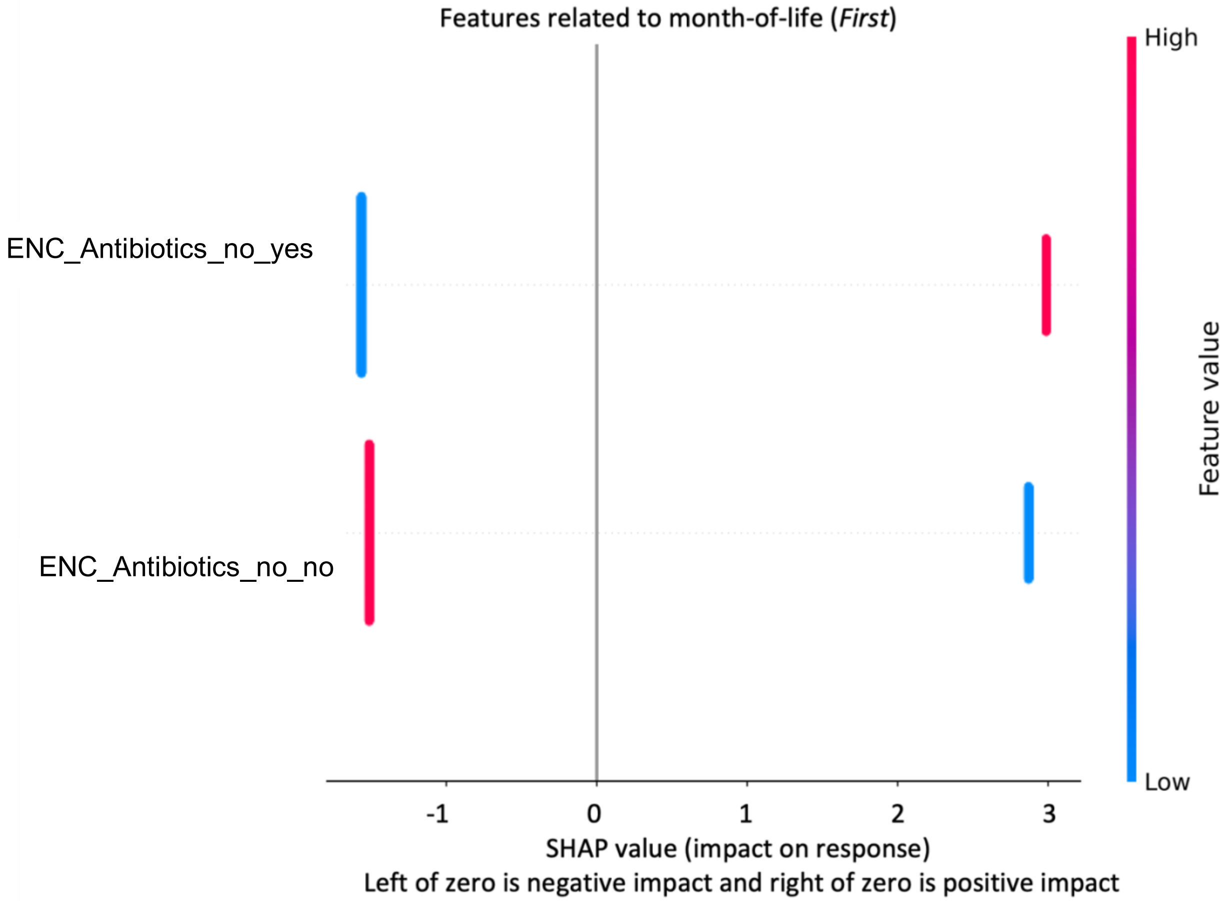


**Supplemental Figure 4. Heatmap of bacterial genera predictive of month-of-life in newborns selected by EXPLANA using only categorical features from the Early Childhood and Microbiome (ECAM) dataset**. RF was used with 500 trees, a feature fraction of 0.2, max depth of 7, and 100 BorutaSHAP trials (100% threshold; p=0.05). Each point represents one sample, and the horizontal position indicates impact on the outcome as indicated on the x-axis. Points to the left indicate a negative impact, and points to the right indicate a positive impact. The colors represent the selected feature values, where red is larger, and blue is smaller. For binary encoded features (‘ENC’) red is yes/1 and blue is no/0.

#### Supplemental Tables

Supplemental Table 1. Engineered predictive and not-predictive input features used in happiness simulation studies.

| **Input variable** | **Predictive or Not Predictive** | **Categorical or Numerical** | **Motivation for inclusion** | **Description of expected effect** |
|---|---|---|---|---|
| Timepoint | Predictive | Numerical (original dataset); Categorical (delta datasets) | Timepoint is needed for longitudinal models and helps explain changes over time for other features | Correlation with changes in happiness over time |
| Cohort | Predictive | Categorical | feature with a positive impact on happiness and no relationship to time | Unhealthy = negative effect; healthy = positive effect |
| Intervention | Predictive | Categorical | has both a time-independent positive impact on happiness and a time-dependent, positive linear relationship to happiness | Treated and therapy_2: positive linear relationship to time with therapy_1 having a bigger impact |
| Pill color | Predictive | Categorical | Categorical variable with feature changes that impact response in an order-dependent manner | green_blue has negative impact (can only be identified in Previous and Pairwise models as green did not occur at baseline) |
| Sunshine | Predictive | Numerical | Numerical variable with positive linear relationship to response | Positive linear relationship |
| Relaxation | Predictive | Numerical | Effect that plateaus and remains constant (for varying patterns of change over time) | T1 has one value; T2, T3, T4, T5 have the same equivalent value (different than T1) |
| Salary | Predictive | Categorical | Categorical variable with two features that impact response | High has a positive impact and low has a negative impact |
| Pet | Predictive | Categorical | Categorical variable with two features that impact response | |
| Random data distributions: normal, Bernoulli, binomial, Poisson, exponential, gamma, Weibull, and Dirichlet | Not Predictive | Numerical and Categorical representation (binary data distribution) | To test workflow performance with random variables that are not predictive. Random variables are included in *SimFeaturesMicrobiome500* and *SimFeaturesMicrobiome1000* | No anticipated relationships |
| Differentially abundant microbes | Predictive | Numerical | To test workflow performance with sparse, compositional simulated microbes that increase over time along with response. Included in *SimMicrobiome* and *SimFeaturesMicrobiome0*, *SimFeaturesMicrobiome500,* and *SimFeaturesMicrobiome1000* | Positive linear relationship to happiness |
| Not differentially abundant microbes | Not Predictive | Numerical | To test workflow performance with compositional data that does not increase over time. Included in *SimMicrobiome* and *SimFeaturesMicrobiome0*, *SimFeaturesMicrobiome500,* and *SimFeaturesMicrobiome1000* | No anticipated relationships |
